# *Toxoplasma* F-Box Protein 1 is Required for Daughter Cell Scaffold Function During Parasite Replication

**DOI:** 10.1101/511386

**Authors:** Carlos Gustavo Baptista, Agnieszka Lis, Bowen Deng, Elisabet Gas-Pascual, Ashley Dittmar, Wade Sigurdson, Christopher M. West, Ira J. Blader

**Affiliations:** Department of Microbiology and Immunology, University at Buffalo School of Medicine, Buffalo, NY 14203; Department of Biochemistry & Molecular Biology, University of Georgia, Athens, GA 30602 USA; Department of Physiology and Biophysics, University at Buffalo School of Medicine, Buffalo, NY 14203

**Author notes:** HarkerBIO, LLC, 700 Ellicott St, Buffalo, NY 14203.

**Keywords:** Cell Cycle, Parasitology, Ubiquitin, Membrane Trafficking, Cytoskeleton

## Abstract

By binding to the adaptor protein SKP1 and serving as substrate receptors for the Skp1, Cullin, F-box E3 ubiquitin ligase complex, F-box proteins regulate critical cellular processes including cell cycle progression and membrane trafficking. While F-box proteins are conserved throughout eukaryotes and are well studied in yeast, plants, and animals, studies in parasitic protozoa are lagging. We have identified eighteen putative F-box proteins in the *Toxoplasma* genome of which four have predicted homologs in *Plasmodium*. Two of the conserved F-box proteins were demonstrated to be important for *Toxoplasma* fitness and here we focus on an F-box protein, named TgFBXO1, because it is the most highly expressed by replicative tachyzoites and was also identified in an interactome screen as a *Toxoplasma* SKP1 binding protein. TgFBXO1 interacts with *Toxoplasma* SKP1 confirming it as a bona fide F-box protein. In interphase parasites, TgFBXO1 is a component of the Inner Membrane Complex (IMC), which is an organelle that underlies the plasma membrane. Early during replication, TgFBXO1 localizes to the developing daughter cell scaffold, which is the site where the daughter cell IMC and microtubules form and extend from. TgFBXO1 localization to the daughter cell scaffold required centrosome duplication but occurred before segregation of the centrocone, which connects the spindle pole to the centrosomes. Daughter cell scaffold localization required TgFBXO1 N-myristoylation and was dependent on the small molecular weight GTPase, TgRab11b. Finally, we demonstrate that TgFBXO1 is required for parasite growth due to its function as a daughter cell scaffold effector. TgFBXO1 is the first F-box protein to be studied in apicomplexan parasites and represents the first protein demonstrated to be important for daughter cell scaffold function.

**AUTHOR SUMMARY:** *Toxoplasma gondii* is a protozoan parasite that can cause devastating and life-threatening disease in immunocompromised patients and in fetuses. Its replication is important to study because parasite growth is responsible for the pathology that develops in toxoplasmosis patients. The parasite replicates by a unique process named endodyogeny in which two daughter parasites develop within the mother cell. Early during this process the parasite creates a structure called the Daughter Cell Scaffold whose function is to mark the site from which daughter parasites will emerge. Here, we report the identification of one of the first proteins recruited to the Daughter Cell Scaffold and its importance in executing its function.

## INTRODUCTION

*Toxoplasma gondii* is an intracellular apicomplexan parasite responsible for one of the most common parasitic infections in humans and animals [1, 2]. Infections are initiated by digesting either bradyzoite-containing tissue cysts or sporozoite-laden oocysts that are disrupted in the stomach [3]. Released parasites then infect the small intestine and convert into tachyzoites, which triggers the recruitment of inflammatory cells, which are in turn infected and used to disseminate throughout the host where they convert into long-lived tissue cysts [4, 5]. Occasionally, cysts reactivate and the released parasites will revert to tachyzoites, which replicate by a unique process termed endodyogeny where the two daughter parasites develop within the mother [6]. Since parasite replication underlies onset of disease in toxoplasmosis patients, it is critical to study the mechanisms regulating endodyogeny.

Formation of the inner membrane complex (IMC) is a critical step in endodyogeny. The IMC is a unique organelle comprised of an intermediate-filament like cytoskeletal network and membranous sacs whose outer leaflet anchors the actin-myosin gliding machinery [7–9] whereas the cytoplasmic leaflet is associated with subpellicular microtubules that extend from the apical end of the parasite for most of its length [10, 11]. During daughter cell development, the IMC emerges from a structure named the daughter cell scaffold (DCS) that is located apically to the parasite’s outer core of its bipartite centrosome [12, 13]. The DCS serves as the docking site for Rab11b-regulated vesicles containing the material needed to form the emerging IMC [14]. Despite its central role in endodyogeny, it is largely unknown what proteins localize to the DCS nor is it understood how the DCS regulates IMC assembly.

F-box proteins are a family of proteins defined by the presence of a F-box domain and can be divided into three classes, FBXW, FBXL and FBXO [15] according to the presence or absence of canonical substrate recognition domains [15, 16]. F-box proteins are best known as subunits of the SKP1/Cullin/F-box containing E3 ubiquitin ligase (SCF-E3) complex [17] in which the F-box domain binds SKP1 to recruit itself and/or other substrates for polyubiquitination [17, 18]. Although well studied in fungi, plants, and metazoans, the function of F-box proteins in apicomplexan parasites remains unexplored. Previously, we reported that an O_2_-regulated *Toxoplasma* prolyl hydroxylase modifies a proline residue in TgSKP1, which can then be modified by a series of glycosyltransferases [19–21]. TgSKP1 prolyl hydroxylation/glycosylation is predicted to alter F-box protein binding leading to changes in substrate recognition [22].

Here, we identify 18 putative F-box proteins in the *Toxoplasma* genome that, relative to other organisms, represents a significantly streamlined repertoire of F-box proteins [23, 24]. We focused on *Toxoplasma* F-box protein 1 (TgFBXO1) and found that it localizes to the DCS early during endodyogeny and is required for DCS function. Thus, TgFBXO1 represents the first protein identified to be an executor of DCS function.

## RESULTS

### Identification of TgFBXO1 as an Apicomplexan Conserved F-box Protein

F-box proteins contain an F-box domain of ∼50 amino acids that are organized into a triple α-helical bundle that mediates binding to SKP1 [25]. We employed a series of reciprocal BLASTp and hidden Markov searches seeded with 43 F-box sequences from known F-box proteins including 7 from crystal structures (Figure S1A) and 37 from two well studied yeast species (Figure S1B), to assemble a list of 18 predicted *Toxoplasma gondii* F-box proteins (Table S1; Figure S1C). Five are of the FBXW and FBXL classes, while the others possess C-terminal domains that are poorly if at all are related to F-box proteins outside of protists (data not shown). All but TgFBXO10 have clear homologs in *Neospora caninum* but only four had clear *Plasmodium* homologs, suggesting rapid evolutionary specialization.

In a parallel study, we sought to identify candidate F-box proteins by investigating the TgSKP1 interactome. To maximize coverage, TgSKP1 was captured i) from lysates of extracellular parasites (80% extracellular) prepared at two different detergent concentrations using bead-bound affinity purified polyclonal antiserum (UOK75) generated against TgSKP1 [19], or ii) by using bead-bound anti-FLAG M2 monoclonal antibody to recover TgSKP1 from detergent lysates of extracellular or intracellular parasites of a strain named TgSKP1^SF^ in which the TgSKP1 locus was modified by the addition of a C-terminal Strep/Flag (SF) tag. To differentiate specific from adventitious interactors, non-immune rabbit IgG was used as a control for the UOK75 immunoprecipitation, and the parental RHΔ*hxgprt*Δ*ku80* strain was used as a control for mAb M2 immunoprecipitation. The immunoprecipitates were subjected to nLC-MS/MS analysis and proteins were assigned from peptide identifications at 1% FDR using Sequest, and quantified based on Abundance in Proteome Discoverer 2.2. After filtering to remove non-specific interactions and major organeller proteins, and relaxing the protein-level FDR for predicted FBPs from the bioinformatics search (Figure. S1), twenty-one proteins were identified as candidate SKP1-interacting proteins. Of these, five were present in both the TgSKP1^SF^ and UOK75 immunoprecipitations (Figures 1A & S2 and Table S2). These included the expected SCF complex proteins TgSKP1, CUL-1 and RBX1, a Skp1 glycosyltransferase (GAT1), and TgFBXO1 (TgGT1_310930). Three additional FBPs predicted from the bioinformatics approach were detected in the TgSKP1^SF^ immunoprecipitates, but two were assigned only with medium confidence.

**Fig 1.**
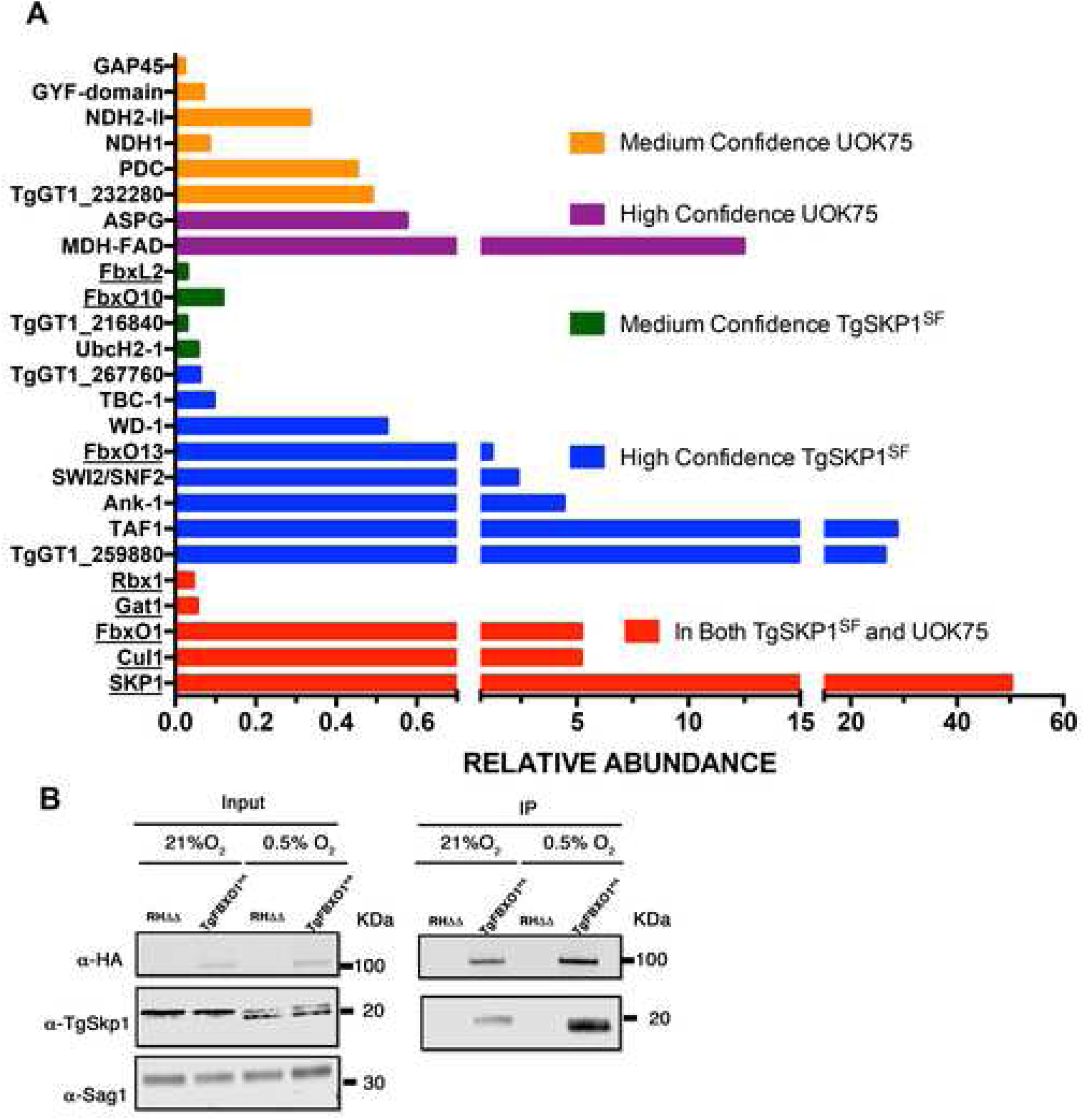
TgFBXO1 Interacts with TgSKP1. **(A).** RHΔhxgprtΔKu80 and TgSKP1^SF^ tachyzoites were solubilized and immunoprecipitated with affinity purified anti-TgSKP1 (UOK75) or anti-FLAG (M2), respectively. The immunoprecipitates were trypsin digested and subjected to proteomic analysis. Protein candidates were selected from a list of protein hits that were detected in multiple replicates and minimally recovered with irrelevant antibodies (for UOK75) or the untagged strain (for M2), and are reported as found in either both (red bars) or only individual immunoprecipitations with either high or medium confidence. Proteins known or predicted to interact with TgSKP1 are underlined. See Figure S2 and Table S2 for more information. **(B).** Lysates prepared from TgFBXO1^HA^ and parental RHΔhxgprtΔKu80 grown at 21% or 0.5% O_2_ were either Western blotted (Input) or incubated with anti-HA antibodies (IP). The immunoprecipitates were captured by Protein G Sepharose and the immune complexes were Western blotted to detect TgFBXO1^HA^, TgSkp1, and SAG1.

We focused efforts on TgFBXO1 because it is conserved in *Plasmodium*, is important for *Toxoplasma* fitness according to a genome-wide CRISPR screen [26], and is the most highly expressed *Toxoplasma* F-box protein throughout the tachyzoite cell cycle https://toxodb.org/toxo/app/record/gene/TGME49_310930. To confirm that TgFBXO1 interacts with TgSKP1, we cloned a C-terminal 3XHA epitope tag onto TgFBXO1 (TgFBXO1^HA^) via homologous recombination [27]. Western blotting lysates from TgFBXO1^HA^ expressing parasites but not the parental RHΔhxgprtΔKu80 strain with anti-HA antibodies revealed a single immunoreactive band at ∼100 kDa, which is the approximate expected molecular weight of TgFBXO1^HA^ (Figure 1B; Input 21% O_2_). Next, TgSKP1 was detected by Western blotting TgFBXO1^HA^ immunoprecipitates with anti-TgSKP1 antisera (Figure 1B; IP 21% O_2_). Taken together, the FBP and interactome discovery pipelines led to the identification of TgFBXO1 as an evolutionarily conserved F-box protein that that stably associates with TgSKP1, and is potentially a major component of the SCF-E3 complex in *Toxoplasma*.

Previously, we reported that ^154^Pro of TgSKP1 is hydroxylated and glycosylated and that these post-translational modifications are predicted to change the conformation of the F-box protein-binding region of TgSKP1 as they do in the social amoeba *Dictyostelium discoideum* when a conserved proline residue in DdSKP1 is similarly modified [22]. An O_2_-regulated prolyl hydroxylase modifies ^154^Pro and under low O_2_ conditions can be observed by Western blotting as a decrease in the apparent molecular weight of TgSKP1 [19, 20, 28] (Figure 1B; TgSKP1 Input Fractions). To test whether TgSKP1 and TgFBXO1 interactions were O_2_-dependent, TgSKP1 levels were compared in TgFBXO1^HA^ immunoprecipitates from parasites grown at either 21% or 0.5% O_2_. The data revealed modestly increased levels of TgSKP1/TgFBXO1 interactions at 0.5% O_2_ (Figure 1B; compare 21% O_2_ to 0.5% O_2_).

### TgFBXO1 is Important for Parasite Growth

Consistent with the finding that it is important for parasite fitness [26], attempts to generate TgFBXO1 knockout by either CRISPR or insertional mutagenesis were unsuccessful (not shown). TgFBXO1 was also resistant to tagging with the relatively large Destabilization Domain [29]. We therefore used a conditional knockdown system where the TgFBXO1 native promoter was replaced with an anhydrotetracycline (ATC)-repressible promoter and an amino-terminal 3XHA epitope tag was cloned in frame to generate ^HA(ATC)^TgFBXO1 (Figure 2A) [30]. ^HA(ATC)^TgFBXO1 protein was undetectable after 24 h treatment with 1 μg/ml ATC (Figure 2B). Next, ^HA(ATC)^TgFBXO1 growth was assessed by plaque assay and found that relative to the parental strain growing with or without ATC that a ∼60% decrease in numbers of plaques were present when ^HA(ATC)^TgFBXO1 expression was decreased (Figures 2C and S2). Moreover, the size of the ^HA(ATC)^TgFBXO1 plaques that did form were ∼70% smaller than plaques formed by ^HA(ATC)^TgFBXO1 expressing parasites (Figures 2D and S2). Although TgFBXO1/TgSKP1 interactions were enhanced at 0.5% O_2_, this growth phenotype was not enhanced at low O_2_ levels. Therefore the rest of this study’s experiments were performed under normoxic (21% O_2_) conditions. Taken together, these data indicate that TgFBXO1 is important for optimal *Toxoplasma* growth.

**Figure 2.**
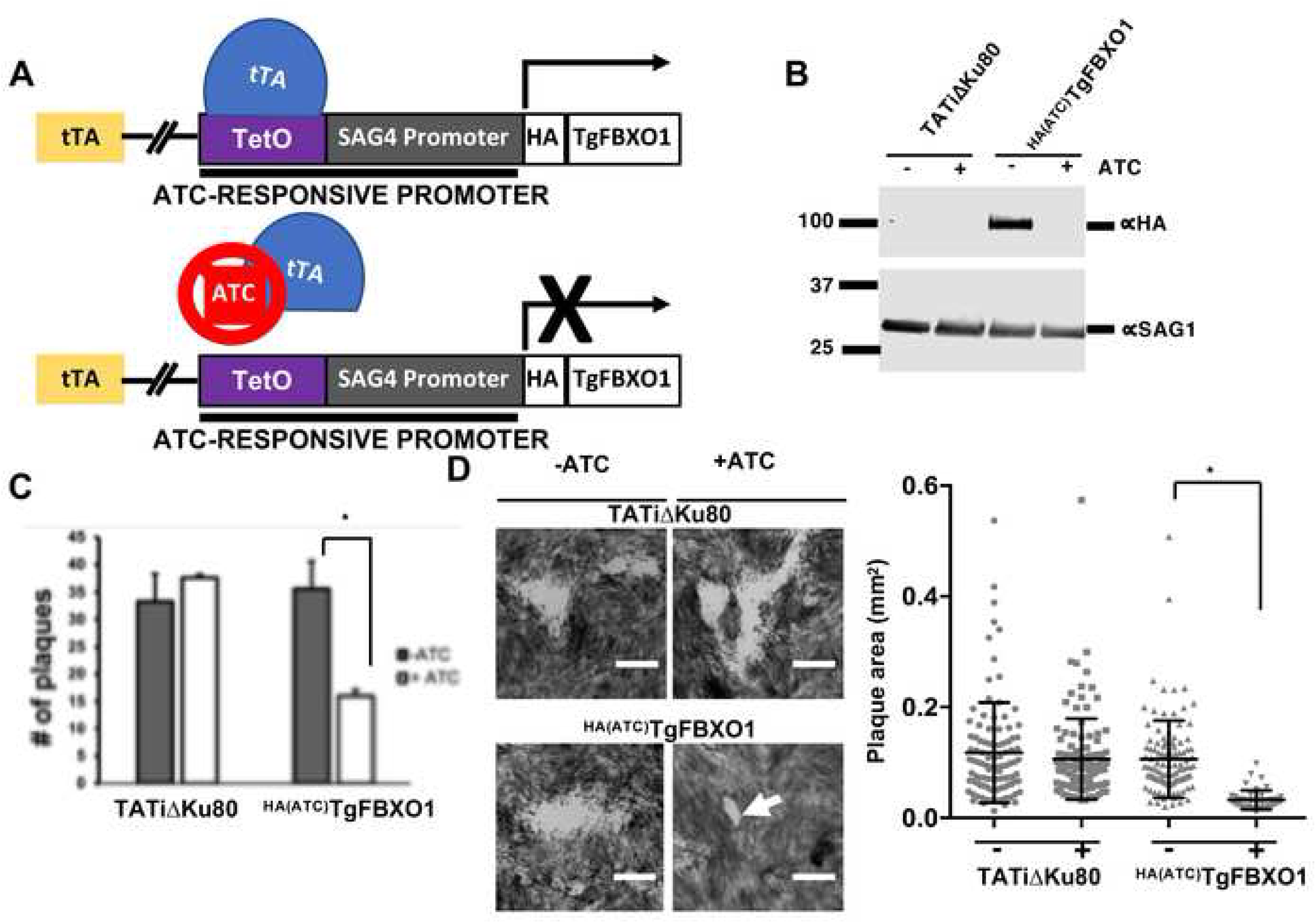
TgFBXO1 is Important for *Toxoplasma* growth. **(A).** Schematic illustration of ^HA(ATC)^TgFBXO1 anhydrotetracycline (ATC)-regulated gene expression. The TgFBXO1 endogenous promoter was replaced with a SAG4 promoter containing a tetracycline transactivator (tTA) binding element. In the absence of ATC, the tetracycline regulated promoter is active, while the addition of ATC reduces transcription by preventing tTA from binding to and activating the promoter. **(B).** Lysates from ^HA(ATC)^TgFBXO1 or parental RHΔhxgprtΔKu80 parasites grown for 24 h in the absence or presence of 1μg/ml ATC were Western blotted to detect ^HA(ATC)^TgFBXO1 or SAG1 as a loading control. **(C).** ^HA(ATC)^TgFBXO1 or RHΔhxgprtΔKu80 parasites (100/well of a 6 well plate) were grown for 5 d on HFF monolayers with or without 1μg/ml ATC. The cells were then fixed and numbers of plaques formed counted. Shown are the averages and standard deviations of 3 independent experiments performed in triplicate. **(D).** Representative images of plaques from (C) showing decreased plaque size of TgFBXO1-depleted parasites. Plaque sizes were quantified and plotted on the adjacent graph. *, p < 0.001, One-Way ANOVA. Bars = 0.5mm.

### TgFBXO1 Forms an Apical Structure Early During Endodyogeny

Although detectable by Western blotting, ^HA(ATC)^TgFBXO1 could not be visualized by immunofluorescence staining suggesting inaccessibility of the N-terminal tag in fixed parasites. We therefore assessed TgFBXO1 localization using the C-terminal-tagged TgFBXO1^HA^ strain and found that in interphase parasites TgFBXO1 localized to the apical end of the parasite and to the periphery where it appeared to colocalize with the IMC protein IMC3, which is a cytoskeletal IMC protein (Figure 3A; G1) [31]. We next examined TgFBXO1 localization during endodyogeny and found that early during endodyogeny TgFBXO1 localized to a perinuclear location before IMC3 associated with daughter IMCs (Figure 3A; S-phase). Later during M-phase (note lobe-shaped nuclei that are migrating into nascent daughter cells), TgFBXO1 remained primarily at the apical end of the growing daughter parasites while IMC3 extended posteriorly (Figure 3A; M-phase).

**Figure 3.**
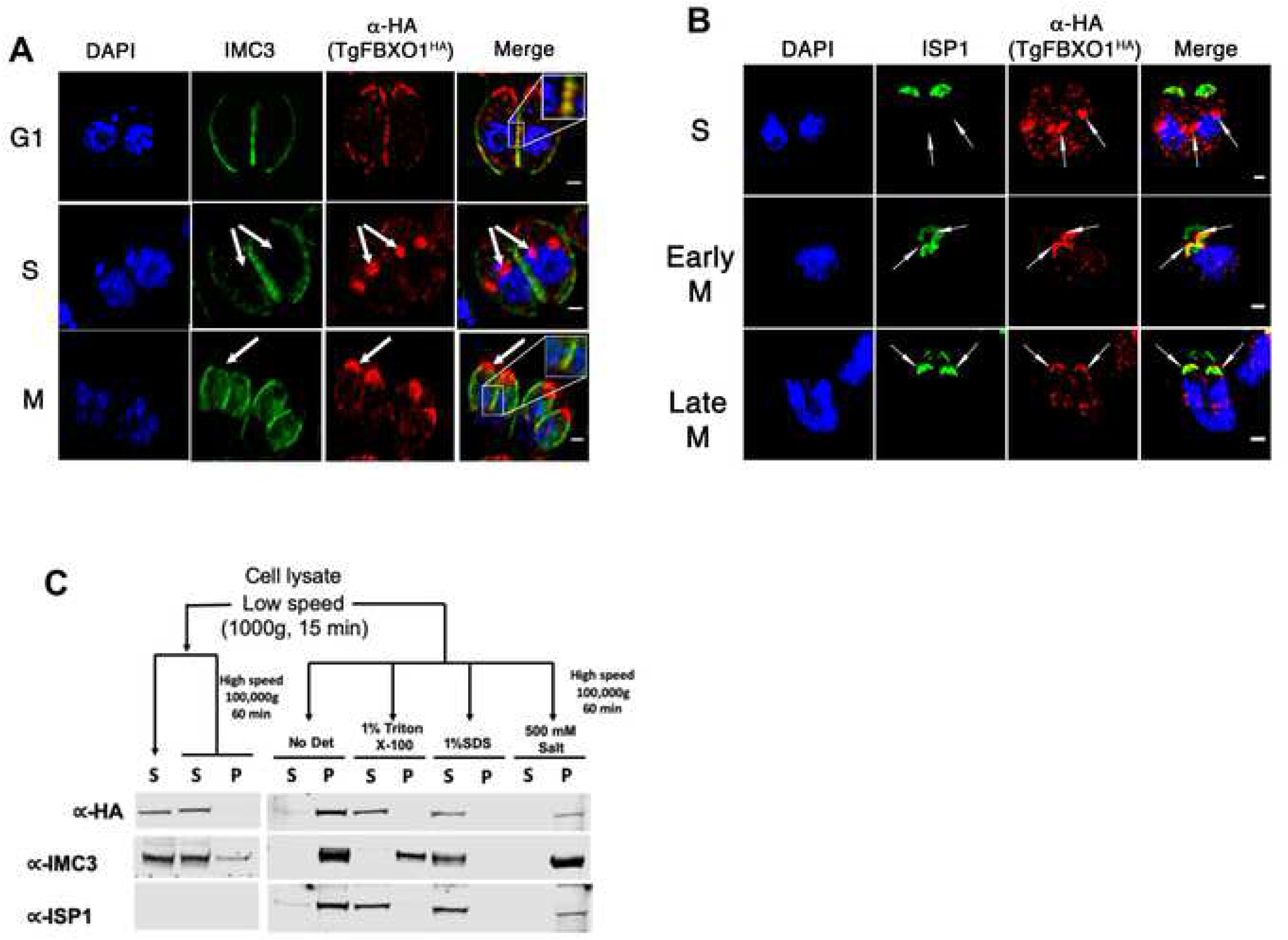
TgFBXO1 Forms Apical Structures Early During Endodyogeny. **(A).** TgFBXO1^HA^-expressing parasites were fixed and stained to detect IMC3, TgFBXO1, and DNA during G1, S, and M phases. Arrows highlight TgFBXO1 apical structures. **(B).** TgFBXO1^HA^-expressing parasites were fixed and stained to detect ISP1, TgFBXO1, and DNA during S, early M (before nuclear segregation), and late M (during nuclear segregation) phases. Arrows highlight TgFBXO1 apical structures. Bars = 2µm. **(C).** TOP: Fractionation scheme to characterize TgFBXO1^HA^ association with the IMC. BOTTOM: Western blotting of equivalent volumes of each indicated fraction. Blots were probed with antibodies to detect TgFBXO1 (α-HA), IMC3 (αIMC3), and ISP1 (αISP1).

ISP1 is another IMC protein that is inserted into the membranous alveoli via fatty acylation. Before IMC3 associates with the daughter IMC, ISP1 forms an apical cap structure in the daughter parasites that remains throughout endodyogeny and into interphase [32]. Comparing ISP1 and TgFBXO1 localization revealed that TgFBXO1 attained its perinuclear localization before ISP1 did (Figure 3B). As parasites progressed through endodyogeny, the ISP1 cap was found near TgFBXO1 although TgFBXO1 was distinct and apical to ISP1 suggesting that the two proteins were in distinct structures. In interphase parasites, apical TgFBXO1 staining did not colocalize with RNG1, which is a component of the parasite’s microtubule organizing center, which is apical to the ISP1 cap (Figure S4) [33], indicating that these two proteins also reside in distinct structures.

We next used biochemical extraction to assess how TgFBXO1 associates with the IMC [34]. Parasites were mechanically lysed under hypo-osmotic conditions and cytoskeletal fractions were collected by low speed centrifugation. The pellets were resuspended in isotonic buffer and IMC embedded proteins were differentially extracted with either high salt (to disrupt electrostatic interactions), Triton X-100 (to extract membrane-associated proteins) or SDS (to extract cytoskeletal proteins). TgFBXO1 was extracted with both Triton and SDS indicating that it is most likely associated with the IMC as a membrane-associated protein in a manner similar to ISP1 (Figure 3C). However, we noted that in contrast to ISP1 that was exclusively associated with the IMC that a significant proportion of TgFBXO1 was soluble.

The first step in endodyogeny is duplication of the parasite’s novel bipartite centrosome followed by duplication of the centrocone (the parasite’s unique spindle compartment), assembly of subpellicular microtubules, formation of the IMC, and finally daughter cell budding and emergence [6, 12, 35]. To more definitively determine when during endodyogeny the TgFBXO1 apical structure forms, TgFBXO1^HA^ parasites were fixed and stained with anti-TgCentrin-1 antibodies to detect the outer centrosome core. We were able to detect parasites where TgFBXO1 remained associated with the mother IMC but containing duplicated outer centrosome cores (Figure 4A; S) indicating that centrosome duplication precedes TgFBXO1 recruitment to the forming daughter parasite. As endodyogeny progressed, TgFBXO1 were positioned apically to the duplicated outer centrosome cores (Figure 4A; EM) and as mitosis progressed (denoted by segregating nuclei), TgCentrin-1 staining was associated with medial segments of TgFBXO1 structures (Figure 4A; LM).

**Figure. 4.**
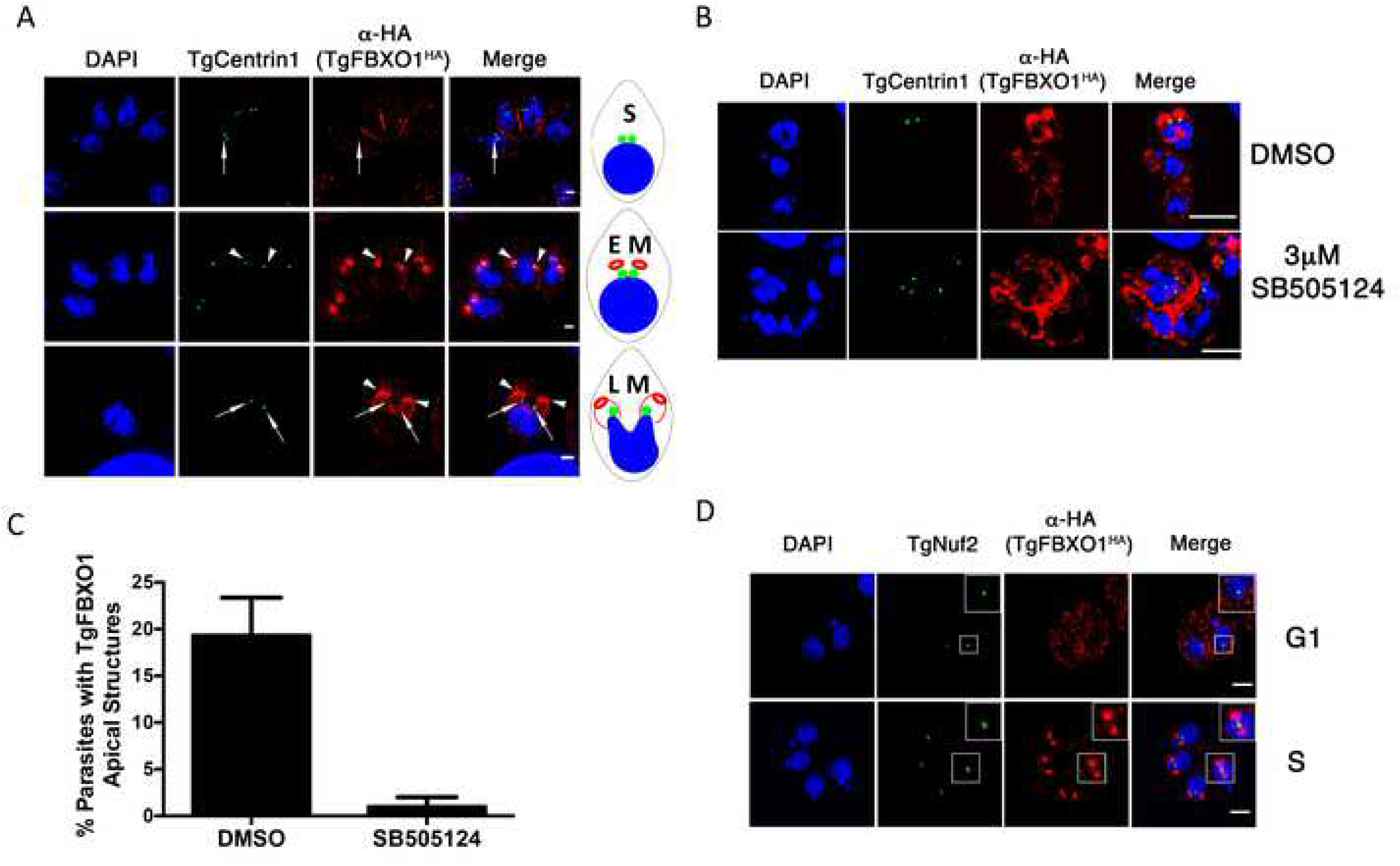
TgFBXO1 Apical Structures Form After Centrosome Duplication and are Localized Apically to the Parasite Centrosome. **(A).** TgFBXO1^HA^-expressing parasites were fixed and stained to detect Centrin1, TgFBXO1, and DNA during S, early M (EM), and late M (LM) phases. Arrows highlight TgFBXO1 apical structures. Bars = 2µm. A schematic depicting the cell cycle phases is shown. **(B).** Parasites were treated with 3 µM SB505124 (or DMSO as a vehicle control) for 24 h, fixed and then stained to detect centrin1, TgFBXO1, and DNA. Note the appearance of SB505124-treated parasites with >2 centrin1^+^ foci and disorganized TgFBXO1 staining. Bars = 5 µm. **(C).** Quantification of numbers of parasites with apical TgFBXO1 structures. Data represents averages and standard deviations of three independent experiments with at least 50 parasites examined/experiment **(D).** TgFBXO1^HA^ parasites were stained to detect TgFBXO1 and the centrocone marker TgNuf2. Note TgFBXO1^HA^ apical structures form after centrocone duplication but before separation as evidenced by the lobed TgNuf2 staining (Insert). Bars: 2µm.

*Toxoplasma* genome duplication and daughter cell development are coordinated by the parasite MAP kinase, TgMAPK-L1, and in the absence of TgMAPK-L1 signaling centrosome duplication continues although IMC development cannot [30]. We therefore tested whether the apical TgFBXO1 structures formed independently of centrosome duplication by treating TgFBXO1^HA^ parasites with either DMSO (vehicle control) or the TgMAPK-L1 inhibitor, SB505124 [36]. In contrast to control parasites, TgFBXO1^HA^ staining remained peripheral in SB505124-treated parasites containing ≥2 outer centrosome cores and numbers of apical TgFBXO1 structures in daughter parasites were significantly reduced (Figure 4B&C). Although it is unclear how TgMAPKL1 signaling is linked to TgFBXO1 apical structure formation, these data indicate that formation of this structure is dependent on the parasite signaling that genome duplication is completed

The *Toxoplasma* centrocone duplicates and separates after centrosome duplication but before ISP and IMC proteins form the daughter cell bud [30, 37]. We therefore compared timing of TgFBXO1 apical structure formation and centrocone duplication/migration by staining TgFBXO1^HA^ cells with antibodies against the centrocone protein TgNuf2 [38]. We identified parasites with distinct TgFBXO1 apical structures that contained duplicated centrocones (note the lobed appearance of TgNuf2 structures) that had not yet separated (Figure 4D; S).

Subpellicular microtubules, which provide positional and structural cues for daughter cell IMC development and organization [11], form after centrosome duplication and centrocone separation [39, 40]. To assess the relationship between the TgFBXO1 apical structures and daughter cell microtubules, TgFBXO1^HA^ cells were stained with antibodies against acetylated tubulin, which denote stable microtubule filaments [41]. Parasites were identified that contained TgFBXO1 apical staining but before acetylated tubulin could be detected in the daughter cells (Figure 5A). As cell cycle progressed, acetylated tubulin colocalized with apical TgFBXO1 although F-box protein staining was consistently more apical.

**Figure 5.**
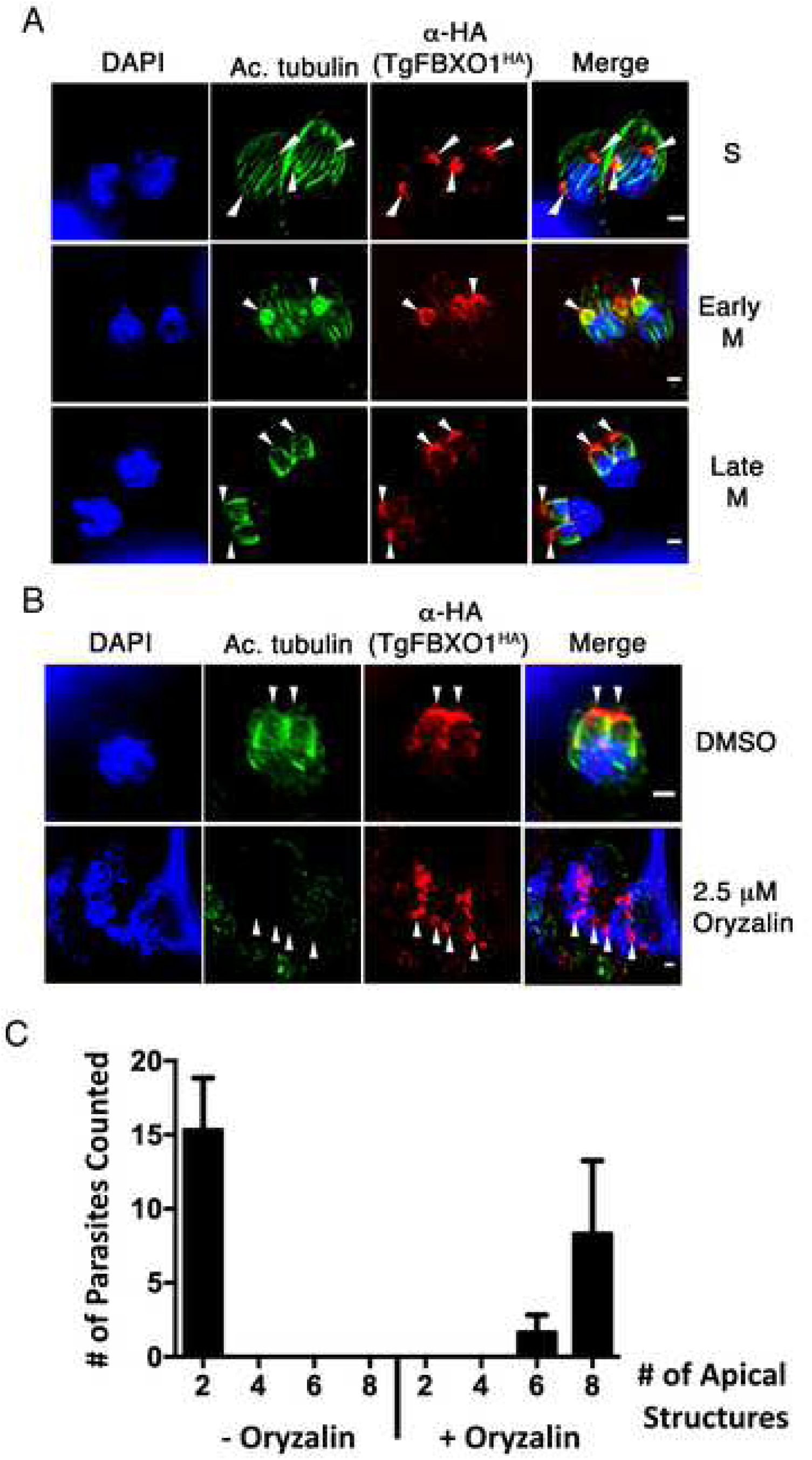
TgFBXO1 Apical Structures form Independently of Daughter Cell Microtubule Assembly. **(A).** TgFBXO1^HA^-expressing parasites were stained to detect TgFBXO1, acetylated tubulin (to detect newly synthesized microtubules), and DNA during S, early M, and late M (during nuclear segregation) phases. Bars = 2 µm. **(B).** Parasites were treated with oryzalin (2.5 µM) or DMSO (vehicle control) for 24 h and then stained to detect TgFBXO1, acetylated tubulin, and DNA. Arrowheads highlight TgFBXO1 apical structures. Bars = 2 µm. **(C).** Quantification of numbers of TgFBXO1^HA^ structures per parasite that form in the presence of DMSO or oryzalin. Shown are the averages and standard deviations of 3 independent experiments in which a total of 50 randomly selected parasites containing TgFBXO1 apical structures were counted.

To test whether TgFBXO1 apical localization was dependent on microtubule assembly, TgFBXO1^HA^ parasites were treated with either DMSO or oryzalin (2.5 μM), which blocks polymerization of parasite but not host microtubules [40]. Oryzalin treatment, which caused the expected defects in microtubule assembly and cytokinesis [42], led to an accumulation of apical TgFBXO1 staining with most parasites possessing 8 structures (Figure 5B&C), indicating that TgFBXO1 apical localization was independent of microtubules.

### TgFBXO1 Is a Component of the Daughter Cell Scaffold

Collectively, the localization data indicate that TgFBXO1 is recruited to a structure that lies apically to the outer centrosome core as well as the emerging IMC. The DCS, which serves as a nucleation site for the daughter IMC and microtubules to emerge from, has such a localization [35, 43]. The small molecular weight GTPase TgRab11b localizes early during endodyogeny to the DCS and regulates IMC protein trafficking to the nascent IMC [14]. To compare TgRab11b and TgFBXO1^HA^ localization, TgFBXO1^HA^ parasites were transfected with a Rab11b expression construct which contains a N-terminal myc-tagged ddFKBP domain to control protein levels by adding a synthetic ligand, Shield1 [14]. Parasites grown in the presence of low Shield1 concentrations (0.1 µM) express low levels of TgRab11b with no apparent effect on growth [14]. Under these conditions, examination of >50 parasites in early S-phase revealed thatTgFBXO1 and TgRab11b co-localized indicating that TgFBXO1 is a DCS component (Figure 6A; S). On the other hand, the two proteins displayed limited colocalization in interphase parasites (Figure 6A; G1).

**Figure 6.**
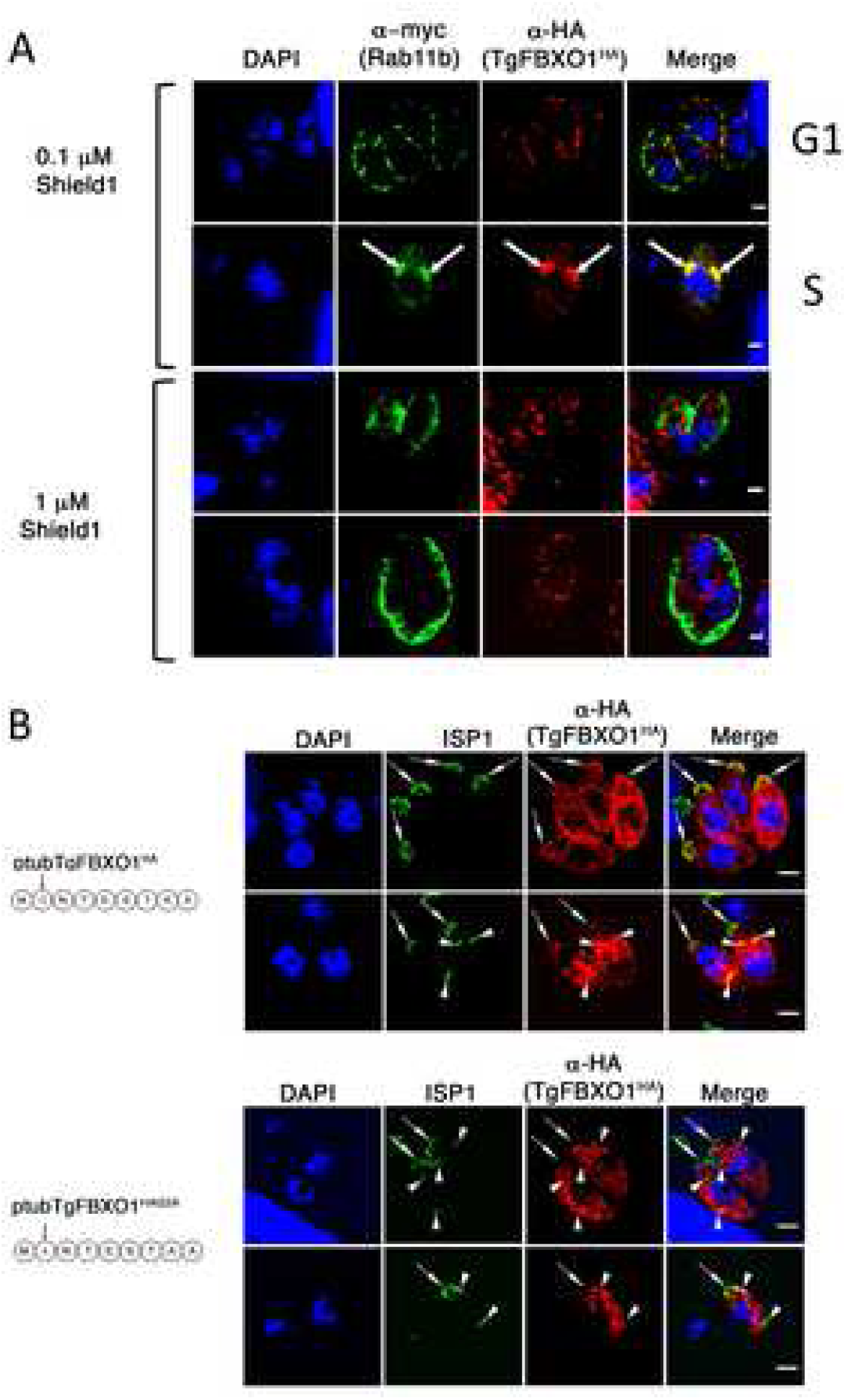
TgFBXO1 Is Recruited to the DCS in a Rab11b-Dependent Manner. **(A).** TgFBXO1^HA^-expressing parasites were transiently transfected with p5RT70DDmycRab11b-HXGPRT expression construct and then grown in the presence of Shield-1 (0.1µM - low level expression or 1 µM – high level expression) reagent for 48 h. Cells were then fixed and stained to detect TgRab11b or TgFBXO1. Arrows highlight apparent colocalization of Rab11b with the TgFBXO1 apical structures. Note peripheral TgRab11b and punctate TgFBXO1 localization in parasites treated with 1 µM Shield-1. Bars = 2µm. **(B).** Parasites were transiently transfected with wild-type (ptubTgFBXO1^HA^) or myristoylation mutant (ptubTgFBXO1^HAG2A^) TgFBXO1 expression constructs, fixed 48 h later, and stained to detect ISP1 and TgFBXO1. Arrows highlight position of IMC1 apical caps. Bars = 2 µm.

To test whether DCS targeting of TgFBXO1 is TgRab11b dependent, TgRab11b-transfected TgFBXO1^HA^ parasites were grown with 1 µM Shield1, which leads to TgRab11b dysfunction due to its overexpression [14]. Under these conditions, overexpressed TgRab11b was mislocalized from the DCS and Golgi to the periphery (Figure 6A 1 bottom panels), which is consistent with [14]. TgFBXO1 was also mislocalized but unlike TgRab11b appeared as cytoplasmic punctae.

The appearance of TgFBXO1 cytoplasmic punctae in TgRab11b overexpressing parasites suggests that TgFBXO1 traffics to the DCS via a membrane trafficking pathway. In addition, TgFBXO1 detergent extractability from the IMC (Figure 3C) suggested that it is membrane-associated. The amino acid sequence of TgFBXO1 predicts neither a transmembrane domain nor a signal sequence but does have a consensus N-myristoylation site at ^2^Gly and this site is conserved amongst the coccidian TgFBXO1 homologs (Figures S5 and S6). To test whether this potential N-myristoylation site is important for DCS and IMC targeting, RHΔhxgprtΔKu80 parasites were transfected to express either wild-type TgFBXO1 (ptubTgFBXO1^HA^) or a mutant where ^2^Gly is mutated to an alanine (ptubTgFBXO1^HAG2A^). In both interphase and mitotic parasites, ptubTgFBXO1^HA^ was properly localized to the periphery (arrow) and DCS (arrow heads), respectively (Figure 6B). In contrast, ptubTgFBXO1^HAG2A^ localization appeared to be largely cytoplasmic and was not localized to either the IMC or DCS.

### TgFBXO1 is Important for DCS Function

To test the importance of TgFBXO1 in DCS function, ^HA(ATC)^TgFBXO1 (or parental TATiΔKu80) parasites were grown in the absence or presence of ATC, fixed 1-5 days later and TgCentrin-1/ISP1 were detected by immunofluorescence staining. In TgFBXO1-depleted daughter parasites, ISP1 was localized near TgCentrin-1 but it was disorganized and the typical apical caps observed in untreated parasites were lacking (Figures 7A and S7A). Development of this phenotype was time dependent and by 5 days post-infection, was evident in ∼76% parasites. Similarly, the ISP1 cap normally found at the mother cell’s apical end was absent in ∼80% of TgFBXO1-depleted parasites by 5 days post-infection (not shown).

**Figure 7.**
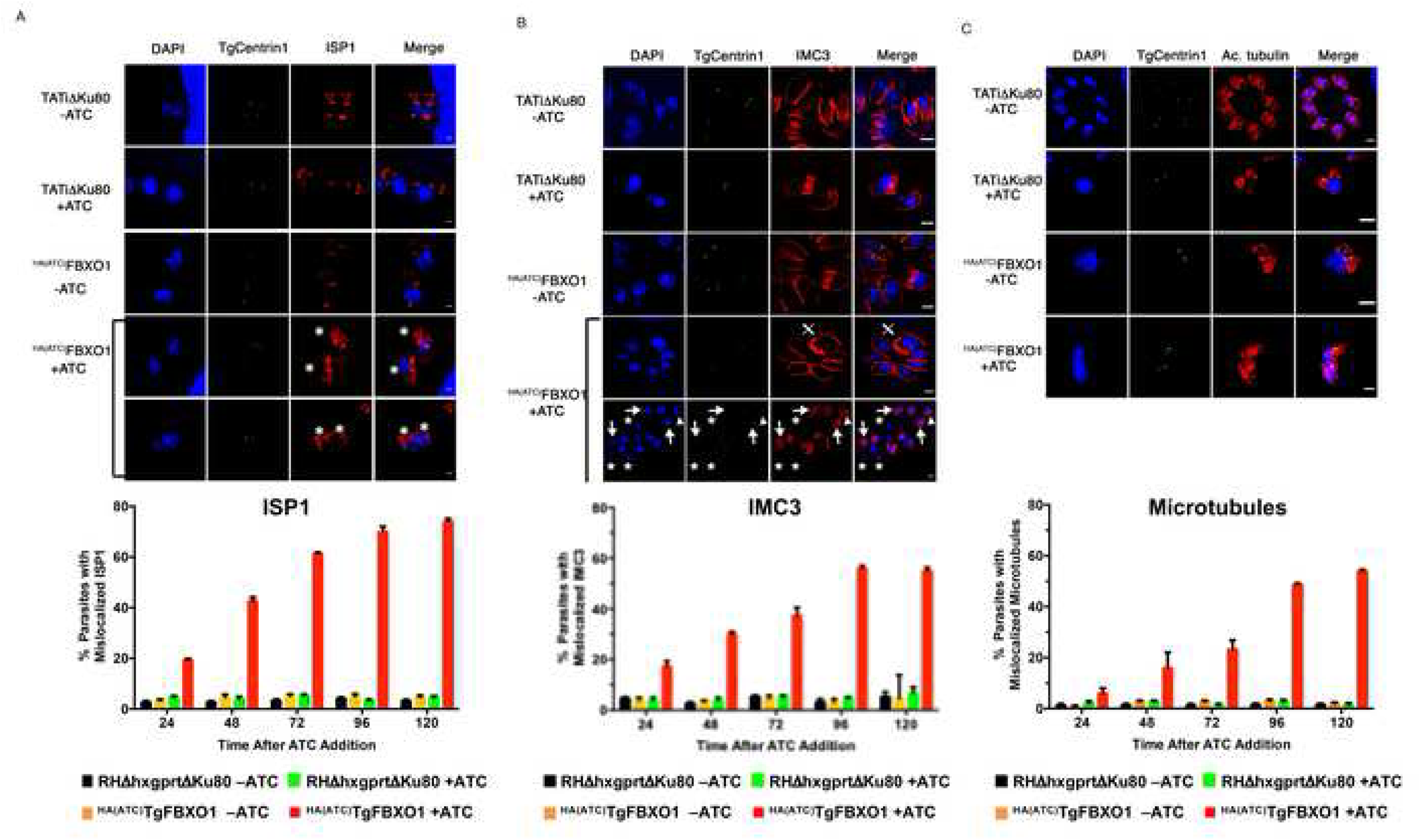
TgFBXO1 Is Required for IMC Organization and Maturation. **(A-C).** Parental RHΔhxgprtΔKu80 and ^HA(ATC)^TgFBXO1 parasites were grown for 1-5 days in the absence or presence of ATC (1 µg/ml) and stained to detect DNA (blue), centrin (green), and ISP1, IMC3, or acetylated tubulin (red). Images are of representative parasites 5 days -/+ ATC. Graphs below each panel represent averages and standard deviations from 3 independent experiments with at least 50 parasites counted per condition. Bars = 2 µm. **(A). \*** denotes mislocalized ISP1 in DCS of parasites with decreased TgFBXO1 expression. Bars = 1 µm. **(B). ⊕** highlights IMC3 whirls, * highlights parasites lacking IMC3 staining, arrows highlight misaligned IMC3 and centrin1, and arrowheads highlight parasites lacking centrin1 staining. Bars = 2 µm. **(C).** Note disorganized appearance of acetylated microtubules in ATC-treated ^HA(ATC)^TgFBXO1 parasites. Bars = 2 µm.

Next, we assessed IMC3 localization and observed a time-dependent increase in IMC3-containing daughter cell buds that were malformed relative to TgCentrin-1. This phenotype was displayed as either: i) IMC3-labelled daughter buds associated with nuclei but centrin-1 staining was absent (Figures 7B (arrows) and S7B). ii) Dividing parasites with only a single IMC (Figures 7B (triangles) and S7B). iii) Nuclei devoid of both TgCentrin1 and IMC3 (Figures 7B (asterisks) and S7B). We quantified numbers of ^HA(ATC)^TgFBXO1-depleted parasites with IMC3 defects and found that, like ISP1, they progressively increased to ∼55% of the parasites counted. Finally, we also noted interphase parasites containing IMC3 whirls (Figures 7B (**X**) and S7B;), which was reminiscent of the effect of TgRab11b overexpression on IMC3 localization [14].

Besides IMC development, the DCS also promotes assembly of the daughter cell subpellicular microtubules. To examine whether microtubule assembly is affected in TgFBXO1-depleted parasites, ^HA(ATC)^TgFBXO1 parasites were grown in the absence or presence of ATC and then newly assembled microtubules were imaged using anti-acetylated tubulin antibodies. In contrast to parasites grown in the absence of ATC, microtubules in ^HA(ATC)^TgFBXO1 parasites with ATC were often missing or not properly organized by five days after adding ATC (Figures 7C and S7C). However, these cytoskeletal defects took longer to develop than either the ISP1 or IMC3 phenotypes suggesting that TgFBXO1 likely regulates IMC development and that the microtubule phenotypes are a secondary consequence.

### Identification of DCS Microdomains

Having established TgFBXO1 as a DCS effector protein, we next used super resolution microscopy to gain insight into the organization and structure of this complex. First, we sought to define the spatial organization between the DCS and the developing IMC by staining dividing TgFBXO1^HA^ parasites with anti-IMC3 and anti-HA antibodies. At an early stage during endodyogeny (after IMC3 begins to accumulate at the daughter cell bud but before it is cleared from the mother cell IMC), TgFBXO1 was found to accumulate as a distinct cap-like structure that was positioned apically from IMC3 structure (Figure 8A; Video S1). However, we also noted that TgFBXO1 extended posteriorly from the cap. In addition, TgFBXO1 was not homogenously distributed throughout the DCS but rather appeared as radial spikes (Figure 8A). TgFBXO1^HA^ expressing parasites were similarly stained to detect TgFBXO1 and ISP1. In contrast to IMC3 whose localization was distinct from TgFBXO1, ISP1 was more proximal to TgFBXO1 although TgFBXO1 was positioned more apical than ISP1 (Figure 8B and Video S2).

**Figure 8.**
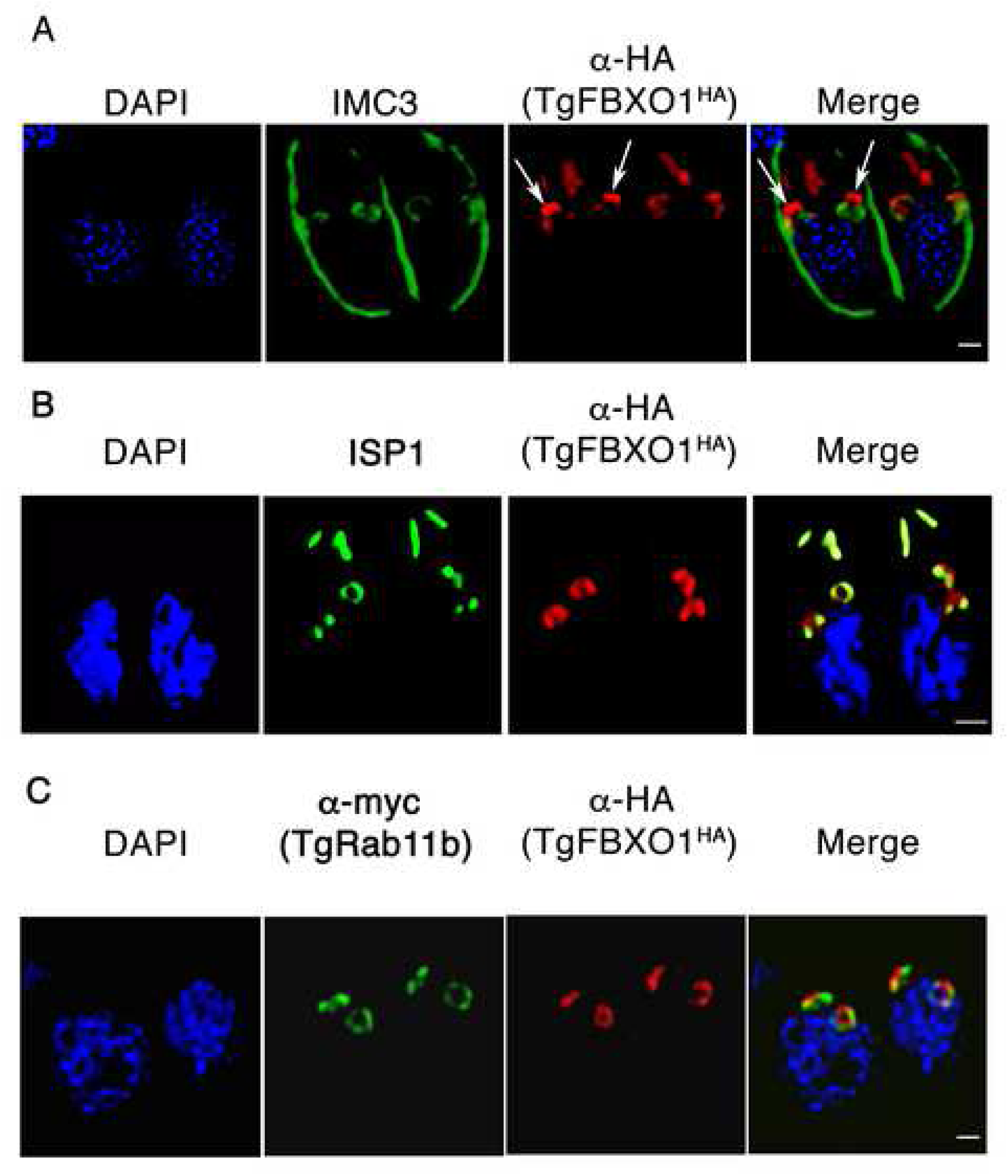
Super Resolution Microscopy Imaging of TgFBXO1. TgFBXO1^HA^ parasites were fixed and imaged using Hyvolution Acquisition Software to detect TgFBXO1 and IMC3 (A), ISP1 (B), and TgRab11b (C). Shown are still images from movies of three dimensional projections available as supplemental data.

As noted earlier, TgRab11b is another DCS-localized protein. Thus, we assessed TgRab11b and TgFBXO1 localization in dividing parasites using super resolution microscopy. Remarkably, TgFBXO1 and TgRab11b localization did not overlap in the DCS but instead appeared to localize to distinct regions and in some cases appeared to be interdigitated although we cannot exclude the possibility that antigen accessibility limits detection of either protein (Figure 8C and Video S3). However, we discount this possibility since examination of > 50 parasites revealed only interdigitation but no colocalization between TgFBXO1 and TgRab11b. Taken together, these data indicate the DCS is likely composed of at least different microdomains and that TgFBXO1 and TgRab11b are distinct markers for each.

## DISCUSSION

Comprising a ubiquitous eukaryotic protein family, F-box proteins play important roles in diverse biological functions [15, 16]. Although extensively studied in metazoans, plants, and fungi, little is known about their function(s) in protists and in particular apicomplexans. Here, we identified 18 putative *Toxoplasma* F-box proteins, which approximates the number found in two well studied yeast systems but is considerably less than the number in *Dictyostelium*, humans and other organisms [23, 24]. TgFBXO1 was reported to be important for parasite fitness [26]; a finding we corroborate using a conditional expression system. We also demonstrated by reciprocal co-immunoprecipitation assays that TgSKP1 and TgFBXO1 interact with each other supporting TgFBXO1 designation as a F-box protein. We further found that TgFBXO1 is a DCS component; a conclusion supported by multiple lines of data. First, it localizes to a structure positioned apically to the daughter cell’s outer centrosome core and developing IMC. Second, daughter cell TgFBXO1 apical structures form before ISP1 and IMC3 recruitment to developing daughter parasites. Third, TgFBXO1 becomes localized to the DCS after centrosome duplication but before centrocone separation. Finally, TgFBXO1 colocalized with TgRab11b at the DCS, albeit in distinct microdomains, and TgFBXO1 recruitment to the DCS was TgRab11b-dependent.

Sequence homology-based studies suggest that proteins related to TgFBXO1 consist of at least 3 domains (Figure S5). The C-terminal domain and features of the middle predicted F-box domain are conserved in proteins throughout the alveolate clade of protists (Figure S5), which includes ciliates and dinoflagellates as well as apicomplexans. The protein acquired a 330-amino acid N-terminal domain only in *Toxoplasma* and closely related cyst-forming coccidia (no trace of this sequence exists in other extant proteins). Thus, a core function of the TgFBXO1 clade may have emerged early in evolution to serve a coccidia specific function, such as organization of the membranous villi that lie underneath the plasma membrane, which is the IMC in apicomplexans [44–46].

Decreased TgFBXO1 expression had distinct effects on two types of IMC proteins. The IMC alveoli meshwork embedded protein IMC3 formed daughter cell IMCs that were not properly aligned with the outer centrosome core. In contrast, ISP1, which is component of a distinct IMC subcompartment, was unable to form its apical cap structure during endodyogeny and this phenotype was maintained in interphase parasites. It should be noted that ISP1 is not an essential gene thus indicating that it is likely that the observed growth defects are not due to ISP1 [32] but rather ISP1 is serving as a marker for the ISP compartment. While these data are consistent with an emerging model in which IMC components are recruited to the daughter cell through distinct mechanisms [32], they also reveal that the DCS is required for both to occur. It is also possible that only ISP1 or IMC3 require TgFBXO1 and that defects observed with the other protein is a downstream effect. Discrimination between these two possibilities awaits characterization of TgFBXO1-interacting proteins.

IMC protein trafficking to the DCS and emerging bud is a poorly understood process. The small molecular weight GTPase, TgRab11b, is required for this process although it remains unclear where it executes its function and what are its effectors [14]. TgRab11b and TgFBXO1 both localize to the DCS but their effects on IMC proteins differed – defects in IMC formation vs misaligned daughter buds, respectively. These differences could be due to different approaches to achieving target protein loss of function (overexpression of an inactive TgRab11b mutant and TgFBXO1 knockdown) or could represent distinct roles for TgRab11b and TgFBXO1 in IMC biogenesis and organization. But given that TgFBXO1 requires TgRab11b for its DCS localization, we favor a model where TgRab11b functions to recruit diverse types of cargo and membrane vesicles to the DCS and that TgFBXO1 is important for executing DCS function in the development of the new IMC. This model is consistent with our finding that the DCS is composed of distinct microdomains and that each likely has distinct functions.

Fatty acylation is emerging as a critical regulator of targeting of IMC and IMC-related proteins to an emerging apicomplexan daughter cells although different proteins utilize different combinations of fatty acid moieties [32, 47–50]. Here, we report that N-myristoylation is required for TgFBXO1 targeting to the DCS. Since N-myristoylation is thought to mediate protein-membrane interactions and is a co-translational process [51, 52], it was surprising that ∼50% of total TgFBXO1 protein was soluble. Explanations for this could include a TgFBXO1-interacting protein that acts to sequester the N-terminal myristoylated glycine or that N-myristoylation of TgFBXO1 occurs post-translationally. While our future work will address these possible mechanisms, these data suggest that TgFBXO1 fatty acylation is a dynamic process that regulates the proteins trafficking and targeting.

Using a conditional expression system, TgFBXO1 was found to be required for optimal parasite growth most likely due to its function in regulating IMC development and organization. TgFBXO1 expression using this system leads to undetectable levels of the tagged protein within 24 h after adding ATC. Yet, the IMC defects were only minimally evident at this time point but rather accumulated with time over the 120 h time course that we performed. One reason for such a delay is that low but undetectable levels of TgFBXO1 are sufficient to sustain replication, a finding previously reported for *Toxoplasma* actin [53]. Alternatively, a second F-box protein can compensate for the loss of TgFBXO1 albeit not as efficiently.

We are currently unable to differentiate whether TgFBXO1 acts in a SCF-E3 ubiquitin ligase dependent or independent manner [15, 54, 55]. It is possible that TgFBXO1 functions in IMC protein ubiquitylation to regulate their turnover and/or targeting during endodyogeny. Such a scenario is supported by the finding that many ubiquitylated *Toxoplasma* proteins are cell cycle regulated including ISP1 and a large number of other IMC proteins [56]. It is also possible that TgFBXO1 is itself an SCF-E3 substrate and its ubiquitylation is required during endodyogeny for either trafficking of TgFBXO1 from the IMC to the DCS or for its turnover from the mature IMC early during endodyogeny. It is noteworthy that a parasite deubiquitinase, TgOTUD3 [15, 54, 55, 57, 58] was recently reported to coordinate mitosis and cytokinesis by properly matching the correct numbers of centrosomes with numbers of daughter buds. However, we do not believe that TgFBXO1 functions in this manner because in contrast to TgOTUD3, loss of TgFBXO1 leads to unmatched numbers of outer centrosome cores and daughter buds without an increase in parasite ploidy.

Alternatively, TgFBXO1 may function in a SCF-E3 ubiquitin ligase independent manner [59–61]. Such a mechanism could occur by the TgSKP1/TgFBXO1 heterodimer, which according to precedent is expected to have high affinity [62–64], being incorporated into a supramolecular complex where TgSKP1 is shielded from interacting with cullin-1 to form a functional SCF complex. Recent cryo-EM studies of the yeast kinetochore provide an example of how the Skp1/Ctf13 (a yeast F-box protein) complex contributes to kinetochore structure in a way where Skp1 is inaccessible to Cul-1 interaction [60, 61]. Certainly, it is conceivable that SCF-E3 dependent or independent mechanisms need not be mutually exclusive, because association with the SCF-E3 could be a mechanism to turn over excess TgFBXO1 that is not associated with the DCS or IMC.

In summary, this work represents the first investigation of F-box proteins in apicomplexan parasites and reveals the first DCS-localized protein that is required for DCS function. Identification of TgFBXO1-interacting proteins will allow us to determine how TgFBXO1 executes this function. In addition, it remains to be determined whether TgFBXO1/TgSKP1 interactions are regulated by O_2_-dependent hydroxylation and glycosylation of TgSKP1 and whether these impact DCS function or another process.

## MATERIALS AND METHODS

### F-box Homology Search Methods

Previously, 4 putative F-box-like motifs were identified in the *Toxoplasma* genome based on a hidden Markov algorithm [65]. Further BLASTp [66] and hidden Markov approaches using F-box-like motifs originating from known F-box domains from 7 crystal structures (Figure S1A) and the entire collection of budding and fission yeast F-box proteins (Figure S1B) identified 14 additional putative F-box proteins in *Toxoplasma* (Figure S1C). Databases searched included EupathD (https://eupathdb.org/eupathdb/) (*Toxoplasma*, *Neospora caninum*, *Plasmodium* spp, and *Eimeria* spp.), *Saccharomyces* Genome Database (https://www.yeastgenome.org), and PomBase (https://www.pombase.org) (*Schizosaccharomyces*). Each candidate was used to re-search the genome until a comprehensive candidate list was created, which was trimmed of sequences that exceeded the range of variation at individual positions of known F-box sequences (see Figures S1A and S1B) with greatest emphasis given to the pattern of hydrophobic residues. Other domain identities were searched using Conserved Domains and Protein Classification search (http://www.ncbi.nlm.nih.gov/Structure/cdd/cdd.shtml) based on database CDDV3.05-42589 PSSMs with E<0.1 [67]. Searches were also performed at Uniprot (www.uniprot.org/).

To identify TgFBXO1 homologues, the NCBI non-redundant database (November 2018) was searched using the standard BLASTp algorithm at e<100, initially with N-, F-, and C-domains (Figure S2) of TgFBXO1 as query sequences. Except for the F-domain, only sequences from alveolates were retrieved. Repeat searches with non-apicomplexan alveolates failed to identify additional sequences outside of alveolates. Top-scoring sequences representing the diversity of known alveolate genomes were initially aligned in COBALT and manually curated for optimal alignment of hydrophobic residues and minimal indels except adjacent to Gly and Pro residues.

### Cell Lines and *Toxoplasma* Strains

All parasite strains, including parental RHΔ*hxgprt*, TgSKP1^SF^ [20], TATiΔ*ku80* [68] and RHΔ*hxgprt*Δ*ku80*) strains [69], were maintained in human foreskin fibroblasts (ATCC; Manassas, VA) in Dulbecco’s Modification of Eagle’s Medium (DMEM) (VWR; Radnor, PA) supplemented with 10% fetal bovine serum (VWR), 2 mM L-glutamine (VWR), and 100 IU/ml penicillin – 100 µg/ml streptomycin (VWR). Parasites were released from host cells by passage through a 27-gauge needle [70]. All parasite strains and host cell lines were routinely tested for Mycoplasma contamination with the MycoAlert Mycoplasma Detection Kit (Lonza, Basel, Switzerland) and found to be negative.

### Endogenous Epitope Tagging

The ^HA(ATC)^TgFBXO1 conditional expression mutant containing an N-terminal 3x-HA tag was generated by amplifying the 5’ end of TgFBXO1 using the following primers (regions of homology to TgFBXO1 are underlined) 5′-GCCAGATCT<u>ATGGGCAACACGGAATCC-</u>3′ and 5′-GCCGCGGCCG<u>CCGATTGATTTTCATGAACCAGTGTG</u> 3′, digested with BglII/NotI and ligated to replace the CEP250 cassette in ptetO7sag4-HA-CEP250-DHFR-TS [30]. The resulting construct was linearized, transfected into the TATiΔku80 strain [30, 68], and clones isolated by limiting dilution using pyrimethamine resistance.

TgFBXO1^HA^, containing a C-terminal 3X-HA tag, was generating by PCR amplifying the 3’ region of TgFBXO1 using primers 5’-TACTTCCAATCCAA<u>TTTAGCTGAAGCAAGTCCCGAAAGAT</u> 3′ AND 5’-TCCTCCACTTCCAATTTTAGC<u>CGCATTGGCTCCGCCCT</u> 3′ and incorporated into plasmid pLIC-HA3X-HXGPRT via ligation independent cloning [69]. The construct was linearized and transfected into RHΔ*hxgprt*Δ*ku80* strain by electroporation and clones isolated by limiting dilution using mycophenolic acid/xanthine resistance.

ptubTgFBXO1^HA^ was generated by synthesizing TgFBXO1 coding sequence (GenScript, Piscataway, NJ) and ligating it into p5RT70DDmycRab11b-HXGPRT [14] to replace the DDmycRab11b cassette. Mutation of glycine at position 2 to alanine was accomplished using the Quick-Change Mutagenesis kit (Agilent, Santa Clara, CA) to generate ptubTgFBXO1^HAG2A^.

### TgSKP1 Interactome Studies

Anti-TgSKP1 (UOK75) was affinity purified using TgSkp1, as described before for *Dictyostelium* Skp1A [28], bound to protein A/G magnetic agarose beads (Pierce, 78609) at 4 mg IgG protein/ml, and stably cross-linked with dimethyl pimelimidate as described [71]. Non-immune rabbit IgG (Jackson ImmunoResearch; West Grove, PA) was coupled in parallel in identical fashion. SDS-PAGE analysis indicated that >90% of the bound IgG was covalently linked. TgSKP1^SF^ was captured using anti-FLAG M2 agarose magnetic beads (Sigma). The minimal volume of beads required to maximally capture TgSKP1 (∼60%) or TgSKP1^SF^ (∼80%) was determined by titration and Western blot analysis (data not shown).

RHΔ*hxgprt*Δ*ku80* or TgSKP1^SF^ (RHΔ*hxgprt*Δ*ku80*;*CAT^+^*) tachyzoites were harvested from infected HFF monolayers by scraping and passage through a 27-gauge needle, centrifuged at 2000 × *g* for 8’ at room temperature, resuspended in phosphate-buffered saline, counted, pelleted, and frozen at −80°C. For TgSKP1^SF^ and control untagged strain co-IPs with mAb M2, lysates were prepared from both intracellular (IC), and extracellular parasites (EC). For the anti-TgSKP1 co-IPs with UOK75 and control non-immune IgG, RHΔ*hxgprt*Δ*ku80* parasites (90% intracellular) were analyzed at two detergent concentrations. Frozen parasite pellets (2-6 × 10^8^) were resuspended on ice in IP buffer (50 mM HEPES-NaOH, pH 7.4 with 0.2, 0.5 or 1% Nonidet P-40), varied (100 or 150 mM) NaCl concentration, and protease inhibitors (1 mM PMSF, 10 µg/ml aprotinin, 10 µg/ml leupeptin). Lysates were centrifuged at 21,000 × *g* for 20 min at 4**°**C, and supernatants (1.2 ×10^8^ cell equivalents) wer incubated with 10 µl of antibody-conjugated beads under slow rotation for 1 h at 4**°**C. Beads were captured in a DynaMag-2 magnet (Life Technologies) the unbound fraction removed according to the manufacturer’s directions. Beads were then washed 3× with corresponding IP buffer, 3× with 10 mM Tris-HCl (pH 7.4), 50 mM NaCl, and once with 50 mM NaCl in water. Bound proteins were eluted twice with 60 µl 133 mM triethylamine (TEA, Sequencing Grade, Pierce, 25108) by incubating for 15’ at 25°C and then immediate neutralization with 40 µl of 0.2 M acetic acid. The eluted fractions were pooled, dried under vacuum and reconstituted in 8 M urea in 10 mM Tris-HCl (pH 7.4). The reconstituted eluates were reduced in 10 mM DTT for 40 min at 25°C and alkylated in 50 mM 2-chloroacetamide for 30’ at 25°C. Samples were then diluted to 2 M urea with 10 mM Tris-HCl (pH 7.4) and digested with 10 µg/ml Trypsin Gold (Mass Spectrometry Grade, Promega; Madison, WI) overnight at 25°C. Trypsin activity was quenched in 1% trifluoroacetic acid (TFA, Pierce) on ice for 15’ and centrifuged at 1,800 × *g* for 15 min at 4**°**C to remove precipitate. Peptides were enriched by adsorption to C18 pipette tips (Bond Elut OMIX C18, Agilent; Sunnyvale, CA) and eluted in 100 µl 50% (v/v) acetonitrile (ACN, Optima^TM^ LC/MS Grade, Thermo Fisher; Waltham, MA), 0.1% (v/v) formic acid (FA, LC-MS Grade, Pierce), followed by 100 µl 75% ACN, 0.1% formic acid. Eluted material was vacuum dried and reconstituted in 40 µl 5% ACN, 0.05% TFA.

4 to 8 µl of the reconstituted peptides were loaded onto an Acclaim PepMap C18 trap column (300 μm, 100 Å) in 2% ACN, 0.05% TFA at 5 μl/min, eluted onto an Acclaim PepMap C18 column (75 μm × 150 mm, 2 μm, 100 Å), and eluted with a linear gradient consisting of 4-90% solvent B (solvent A: 0.1% FA; solvent B: 90% ACN, 0.08% FA) over 180 min at a flow rate of 300 nl/min on an Ultimate 3000 RSLCnano UHPLC system, and eluted into the ion source of an Orbitrap QE+ mass spectrometer (Thermo Fisher). The spray voltage was set to 1.9 kV and the temperature of the heated capillary was 280°C. Full MS scans were acquired from m/z 350 to 2000 at 70k resolution, and MS^2^ scans following higher energy collision-induced dissociation (HCD, 30) were collected for the Top10 most intense ions, with a 30” dynamic exclusion. Intervening blank runs ensured absence of carryover between replicates. The acquired raw spectra were analyzed using Sequest HT (Proteome Discoverer 2.2, Thermo Fisher) with a full MS peptide tolerance of 10 ppm and MS^2^ peptide fragment tolerance of 0.02 Da, and filtered to generate a 1% target decoy peptide-spectrum match (PSM) false discovery rate for protein assignments, which were filtered at 1% FDR or relaxed at the protein level to 5% FDR (as indicated). Variable modifications were Met oxidation, Q and N deamidation, and N-terminal acetylation; static modification was Cys carbamidomethylation.

### Plaque Assays

One hundred parasites were added to each well of a 6 well plate containing confluent HFFs in the presence or absence of 1μg/ml ATC. After 5 days, monolayers were methanol-fixed, stained with 0.1% crystal violet, and counted as described [72]. Briefly, plaques imaged using an Olympus SZ61 stereomicroscope equipped with a video camera. Plaque areas were measured using ImageJ software (https://imagej.nih.gov/ij/).

### Immunoprecipitation

TgFBXO1^HA^ tachyzoites grown at 21% or 0.5% O_2_ (in a Baker Hypoxia Chamber (Sanford, ME) were harvested by syringe lysis and washed in ice-cold PBS. Parasites (1×10^8^) were resuspended in 50 mM Tris-HCl pH 7.4 with 1% Triton X-100, 100 mM NaCl, 1 mM NaF, 0.5 mM EDTA, 0.2 mM Na3VO4, 1X protease inhibitor cocktail (Thermo Fisher Scientific), incubated on ice for 30 min and then subjected to sonication. Lysates were clarified by centrifugation at 16,000 *x*g, incubated with mouse α-anti-HA clone 12CA5 conjugated with protein G beads (Sigma-Aldrich) for 16 hours at 4°C. Immune complexes were separated with SDS-PAGE and then Western blotted with antibodies against rat anti-HA clone 3F10 (Roche) or rabbit anti-TgSKP1 UOK75 [19]

### IMC Fractionation

IMC fractionation was done essentially as described in [34]. Briefly, freshly harvested tachyzoites were resuspended in hypotonic lysis buffer (10 mM Tris HCl pH 7.8, 5 mM NaCl) and lysed by repeated freeze thawing followed by centrifugation at 1,000 x*g* to generate fractions S1 (supernatant) and P1 (IMC). S1 was collected and centrifuged at 100,000 x*g* and fractions S2 (cytosol) and P2 (non-IMC membrane) were collected. Fraction P1 was resuspended in isotonic buffer (10 mM Tris HCl pH 7.8, 150 mM NaCl). SDS or Triton X-100 in isotonic buffer were added to P1 to final concentration of 1% and then centrifuged at 16,000 x*g*. Alternatively, P1 fraction in the presence of 500 mM NaCl were centrifuged at 100,000 x*g* for 1 h to isolate proteins that interact with the IMC via electrostatic interactions. All manipulations were performed at 4°C and all buffers were supplemented with 1X Protease Inhibitor cocktail.

### Immunofluorescence Microscopy

*Toxoplasma*-infected HFFs grown on coverslips were fixed either with 4% w/v paraformaldehyde in phosphate buffered saline (PBS) for 20 min at room temperature or by ice-cold methanol for 15 min. Cells were permeabilized with 0.1% Triton X-100 in PBS for 10 min, blocked in 5% w/v bovine serum albumin (BSA) in PBS for 60 min, incubated overnight with primary antibodies (Supplemental Table S2) at 4°C. Coverslips were washed with PBS and then incubated for 60 ‘with Alexa Fluor 488- or Alexa Fluor 594-conjugated secondary antibodies (1∶2000, Thermo Fisher Scientific). DNA was stained either by incubation with 1 µg/ml DAPI (Thermo Fisher Scientific) for 5 minutes followed by mounting in ProLong Glass Antifade Mountant (Thermo Fisher Scientific) or with DAPI-containing VECTASHIELD mounting medium (Vector Labs; Burlingame, CA). Images were acquired using a 100X Plan Apo oil immersion 1.46 numerical aperture lens on a motorized Zeiss Axioimager M2 microscope equipped with an Orca ER charge-coupled-device (CCD) camera (Hamamatsu, Bridgewater, NJ). Images were collected as a 0.2-μm z-increment serial image stacks, processed using Volocity (version 6.1, Acquisition Module (Improvision Inc., Lexington, MA)). Images were deconvolved by a constrained iterative algorithm, pseudocolored, and merged using the Volocity Restoration Module.

For super resolution microscopy, 20 images were acquired as 0.15 µM z-increment serial images using Leica Hyvolution Acquisition Software (Leica; Buffalo Grove, IL) with a Leica TCS SP8 confocal microscope equipped with a 100x/1.47 TIRF oil immersion objective lens and both white light laser (470nm-670nm) and 405 nm diode laser. Image datasets were then deconvolved using Huygens deconvolution software (Hilversum, Netherlands) and 3D volumes generated Leica visualization software. Images from same experiments were processed using identical settings. Unless otherwise noted, data were quantified from at least 50 randomly selected images for each condition from three independently performed experiments.

### Statistical Analyses

When appropriate, one-way ANOVA with Tukey’s post hoc test or Student’s t test was performed with GraphPad Prism (GraphPad, La Jolla, CA).

## ACKNOWLEDGEMENTS

We would like to thank Drs. Peter Bradley, Vern Carruthers, Marc Jan Gubbels, Markus Meissner, Naomi Morrissette, and Elena Suvorova for providing reagents and/or valuable discussion, Kazi Rahman for assistance in predicting F-box domains, and Peng Zhao, Lance Wells, and Steve Hartson for advice on the proteomic studies.

## SUPPORTING INFORMATION LEGENDS

**Figure S1. Known and Predicted F-box Proteins. Figure S1. Known and Predicted F-box Proteins.** F-box domain sequences were predicted as described in Methods. To facilitate visualization of relatedness, acidic residues are in blue, basic in dark red, small or prolines in red, and hydrophobic in green. Positions matching the consensus motif are highlighted in yellow (hydrophobic), gray (acidic), green (basic), or teal (small). F-box sequences from known structures and other validated studies, and functional and interactome studies in *Schizosaccharomyces pombe* and *Saccharomyces cerevisiae*, were aligned to characterize the known diversity of F-box domains. The family of consensus sequences was used to search the genomes of *Toxoplasma* and other apicomplexans. **(A.)** Representative F-box motif sequences derived from crystal structures of F-box protein/Skp1 complexes and other well-known F-box proteins. **(B).** F-box motif sequences from the yeasts *S. pombe* and *S. cerevisiae*. **(C).** Candidate F-box motif sequences identified in the type II ME49 *Toxoplasma* strain.

**Figure S2: Skp1 interactome summary. (A-C)** Intracellular (IC) or extracellular (EC) forms of RHΔhxgprtΔKu80 (RHΔΔ) or Skp1-SF/RHΔΔ tachyzoites were lysed in 0.5% NP-40 and the supernatants were incubated with magnetic anti-FLAG (M2)-beads. The captured material was eluted with 0.15 NH4OH, reduced and alkylated, trypsinized, and analyzed by nLC-MS^2^ in an Orbitrap mass spectrometer. **(A)** Quantitation based on abundance values calculated by the ‘area under the peak’ algorithm in Proteome Discoverer 2.2 from 8 µl sample injections. All proteins that were identified based on 2 or more peptides at 1% FDR, and were not predicted to reside within organelles, and were enriched ≥8-fold in Skp1-SF/RHΔΔ extracts relative to RHΔΔ extracts, are shown. Additional proteins that were seen when criteria were reduced to one peptide and the protein FDR increased to 5%, are included if they were predicted to be in the Skp1 interactome based on F-box sequence motifs or other information. These are labeled in green, and in parentheses if seen only with the relaxed criteria. Proteins whose interaction were supported by detection with a different antibody in panel D, or were observed in a previous unpublished pilot study, are underlined in bold. **(B)** Same as in panel A, from a 4 µl injection. Assignments with higher confidence and at greater abundance were confirmed. **(C)** Comparison of protein abundance, as detected in panel A (8 µl injections), in SF-tagged and untagged extracts. At the right are data from arbitrarily selected examples that were present in both samples, indicating non-specific interactions. The average total Abundance was calculated for this protein set for each sample, and used to generate a normalization factor relative to the Skp1-SF IC values, which was applied to each set of values in Panels A-C. **(D, E)** Similar analysis in which RHΔΔ tachyzoites (80% EC) were lysed in 1% or 0.2% NP-40 and incubated with affinity purified anti-Skp1 (UOK75) or non-specific IgG on magnetic beads. **(D)** Abundance values are reported for all 4 conditions for proteins whose enrichment in UOK75 relative to non-specific IgG was ≥4. **(E)** Values for non-specifically bound proteins. No normalization was performed for this set. See Table S2 for the values and data on the identification criteria.

**Figure S3: Loss of TgFBXO1 Leads to Decreased Plaque Sizes.** Shown are lower magnification images of representative plaques from Figure 2D.

**Figure S4: TgFBXO1 Is Not Associated with the Apical Microtubule Organizing Center in Interphase Parasites.** TgFBXO1^HA^-expressing parasites were transfected with a RNG1-YFP expression plasmid. The parasites were then fixed and TgFBXO1 (αHA; stained to detect TgFBXO1^HA^ (Red), IMC3 (αIMC3; magenta), RNG1 (green), and DNA (DAPI; blue) were detected by Super Resolution Microscopy. Note lack of apparent colocalization between RNG1 and TgFBXO1.

**Figure S5. TgFBXO1 Evolution.** Sequences homologous to TgFBXO1 were aligned in COBALT and subject to manual refinement. To facilitate visualization of relatedness, acidic residues are in blue, basic in dark red, small or prolines in red, and hydrophobic in green. Positions of similarity in one group are highlighted in yellow (hydrophobic), gray (acidic), green (basic), or teal (small). Underlines separate phylogenetic groups: the top two groups consist of different coccidian subclades, the next two groups represent other apicomplexans, the next group contains a chromerid (apicomplexan predecessor), and the bottom group contains non-apicomplexan alveolates including ciliates and dinoflagellates.

**Figure S6. TgFBXO1 Domain Prediction.** Sequence-based homology (Figure S2) suggests that TgFBXO1 consists of at least 3 domains. **(A).** The three domains are conserved in FBXO1 homologues in Coccidians. The N-domain is conserved in length and general amino acid composition, but only the regions shaded in blue, including the N-terminal N-myristylation motif (GxxxS), are conserved in sequence. **(B).** In non-coccidian apicomplexans and other alveolates, the ∼370 C-terminal region is generally conserved. This region includes an N-terminally positioned predicted F-box domain (F) and the C-domain. Some alveolate sequences consist of only F- and C-domains, whereas others contain N-terminal extensions of variable length (dashed line) that bear no detectable sequence similarity to the N-domain of TgFBXO1.

**Figures S7. High Magnification Images of IMC3/ISP1/Acetylated Tubulin Localization in TgFBXO1-Depleted Parasites.** Shown are zoomed in images of each phenotype of the parasites shown in Figures 7.

**Video S1. Animation of 3-D Reconstruction of IMC3/TgFBXO1 Localization.**

**Video S2. Animation of 3-D Reconstruction of ISP1/TgFBXO1 Localization.**

**Video S2. Animation of 3-D Reconstruction of TgRab11b/TgFBXO1 Localization.**

**Supplemental Table S1. *Toxoplasma* F-box Proteins.** Shown are the 18 predicted *Toxoplasma* F-box proteins identified in this study. The ToxoDB or PlasmoDB IDs contain hyperlinks to relevant web pages at the EUPATHDB Genome Database (www.eupathDB.org). Sidik et al Fitness Scores are from [26].

**Supplemental Table S2: List of Candidate TgSKP1 Proteins Identified in the TgSKP1^SF^ and UOK75 Immunoprecipitations**

**Supplemental Table S3: List of Antibodies Used in this Study**

