## Supplemental Figure 1 for "*Toxoplasma* F-Box Protein 1 is Required for Daughter Cell Scaffold Function During Parasite Replication" (from another version of this preprint: version 1, 2019-01-03)

**Fig. S1. Known and predicted F-box proteins.**

**A. Representative F-box motif sequences**

Protein^a^	Identifier	Length Start	Sequence^b^	Other

domain^c^

HsSkp2	GI:21260543	L= 424	94 WDsL**P**D**E**LLLGIFscL-cLPELLKVsGV-cKRWYRLasD--EsLWqtL LRR°

PDB_2ASS	hhhhhhhhhh	hhhhhhhhh	hhhhh	bb^d^

```
MmFbw7         GI:44887884   L= 629  200 IsLLPKELaLYVLsFL-EPKDLLqaaqt-cRYWRILaED--nLLWREK WD40
                 PDB_2OVP                    hhhhhhhhhh   hhhhhhhhhh  hhhhhhhhh
```

HsβTrCP1/Fbw1a gb|AAF04464.1 L= 569 149 pargldhiaenilsyl-dakslcaaelv-ckewyrvtsd--gmlwkkl WD40

```
                 PDB_1P22                    hhhhhhhhhh   hhhhhhhhhhh  hhhhhhhhh    hhhhh
MmFbs1/Fbx02   GI:51315949   L= 297   51 LaELPEPLLLRVLaEL-PatELVqacRLVcLRwKELVDG--aPLwLLK CRD
                 PDB_2E31                      hhhhhhhhh  hhhhhhhhhhh  hhhhhhhh    hhhhhh
AtTir1         GI:332646898  L= 594    5 alsfpeevlehvfsfiqldkdrnsvslv-ckswyeierwcrrkvfign amn1
                 PDB_2P1N                    hhhhhhhhhh    hhhhhhhh    hhhhhhhhhhh bbbb
ScCdc4         GI:14318513   L= 779  275 ItsLPFEIsLKIFnYL-qFEDIInsLGV-sqnwnKIIRK-stsLwKKL WD40
                 PDB_1NEX                    hhhhhhhhhh   hhhhhhhhh   hhhhhhhhhh
ScCtf13        Ymr094w/      L= 478    9 PSFNP-VRFLEL--PIDIRKEVYFHLD@-YAFWLRYDCLV-------- LRR
                 PDB_6FE8                      hhhhhh   hhhhhhhhhhhh   hhhhhhhhhh
ScRcy1         GI:6322257    L= 840    3 DLLKVPEIVTNIASYL-STVDYLSFQQVNKRVYAIINGKNDSKYWSLK
SpPof1p        GI:5913946    L= 605  110 LSLLPVEISFRILSFL-DARSLCQAAQV-SKHWKELADD--DVIWHRM WD40
SpPoplp        GI:162312293  L= 775  301 LTGFPAEITNLVLTHL-DAPSLCAVSQV-SHHWYKLVSS-NEELWKSL WD40
SpPof9p        GI:19112561   L= 467    6 FLELSYDILLEISTYL-DYKDIVHLSET-CKSLSYVFDD--KTIWHRF
HsJHDM1b/Fbl10 GI:54112382   L=1336 1062 aHVMHREVWMaVFsYL-sHqDLcVcMRV-cRtWnRWccD--KRLWtRI JHDM
DdFbxA         DDB_G0276887  L=1247  634 FDNLPEEVVQIIFSNL-SAINIVNLSLV-CKRFKMATDS--PILWKNL WD40
DdFbxD         DDB_G0292312  L= 530  114 ILNLPSTIYAQIFSEF-PVKEILKFSLV-CKEFNKAINH--KFLWKIK WD40
LpAnkB         GI:53751810   L= 154    6 fsdlpeetivntlsfl-kantlgriaqt-cqffnrland+krelwQNL ankyrin
F-box motif consensus sequence*          #xxLPxEILxxILs#L-xxxDLLxLsxV-CbxWxxLsxx--xxIWbxl
```

^a^ Hs, *Homo sapiens*; Mm, *Mus musculus*; At, *Arabidopsis thaliana*; Sc, *Saccharomyces cerevisiae*; Sp, *Schizosaccharomyces pombe*;

Dd, *D. discoideum*; Lp, *Legionella pneumoniae*

^b^ Acidic residues are in blue, basic in dark red, small in red, and hydrophobic in green. Positions matching the consensus motif are highlighted in yellow (hydrophobic), gray (acidic), green (basic), or teal (small). At one position, two alternatives are highlighted. Bolded residues correspond to the PxE motif,

^c^ predominant C-terminal domain type: LRR= leucine rich repeat, WD40= WD40 repeat, amn1= antagonist of mitotic network (LRR-like), CRD= carbohydrate recognition, JHDM= Lys-specific demethylase 2B

^d^ h= α-helical; b= β-strand

+ deleted sequence = khlelhqlrqqri

@=76 aa deleted

* #=hydrophobic side chain, s=small, b=basic, PxE motif is italicized.

**B. F-box motif sequences from the yeasts *S. pombe* and *S. cerevisiae***

Gene name	Length Start Sequence^#^	domains

```
S. pombe
SpPof1p           L= 605  110 LSLLPVEISFRILSFL-DARSLCQAAQV-SKHWKELADD--DVIWHRM WD40
SpPof10p          L= 662   30 VLNLPKEILIIIFSFL-DPRSLLSAQCT-CKYWKKLLSD--DLSWRTA WD40
SpPofllp          L= 506   73 TEVFPEEVSLRVFSYL-DQLDLCKCKLM-SKRWKRLLED--PGIWKAL WD40
SpPoplp           L= 775  301 LTGFPAEITNLVLTHL-DAPSLCAVSQV-SHHWYKLVSS-NEELWKSL WD40
SpPop2p           L= 703  239 LSNLPFSIVQSILLNL-DIHSFLSCRLV-SPTWNRILDV-HTSYWKHM WD40
SpPof2p           L= 463    1  MRVPNEVCFNILSYL-EADELRCKSTV-CTSWRNFII---PTLWEKV LRR
SpPof3p           L= 577  136 FRILPREVLLCILQQL-NFKSIVQCMQV-CKHWRDCIKK-EPSLFCCL TPR/LRR
SpPof7p           L= 361  112 ILKLPDEVLLVILENC+DLRYLSSIALT-CKHFAKALR--ADSLYRSF
SpPof5p           L= 348    1 MQSFPPEIWHHIFDHL-ISFDKFEAKNF-GGLLRICRSSYVGGLHAIY
SpApb1A10(Pof15)  L= 243   31 STLLPVEVIDSVMQYL-PAHDVIQSSFA-SYPLTLIAN----KIIRAR
SpFbhlp           L= 878   11 FYRLPLEIIPLICRFL-SVQDIQSFIRV-FPSFQTILDSSNDLFWKKK
SpPof9p           L= 467    6 FLELSYDILLEISTYL-DYKDIVHLSET-CKSLSYVFDD--KTIWHRF
SpPof8p           L= 402   63 IEPLNPDFLSAVDSIL-EIYFHRERQKE-KVHLAFLIQQ--DDFWKGI
SpPof12p          L= 440   11 ASIFSHETLLHVLNDL-SAHDLAALERV-SRSWNSIVRR--SSVWHNL
SpPof13p          L= 396   44 LFLLNRDIWSLIINYL-DAFDILRLMHS-SRQFYYWLRK--SAVDECC
SpPof6p           L= 872   33 FGcLTINIYLKIFTLI-STPDLCNCRLV-CRKFQQLCDY--NSIYVKK
```

Residues deleted for the alignment: +=IRDLH

```
S. cerevisiae
ScYil046w=Met30   L= 640  184 ISILPQELSLKILSYL-DCQSLCNATRV-CRKWQKLADD--DRVWYHM WD40
ScYfl009c=Cdc4    L= 779  275 ITSLPFEISLKIFNYL-QFEDIINSLGV-SQNWNKIIRK-STSLWKKL WD40
ScYjr090c=Grr1    L=1151  317 LNMLPSEILHLILDKLNQKYDIVKFLTV-SKLWAEIIVK--ILYYRPH LRR
ScYdr306c         L= 478  112 KMVLPWEIQHRIIHYL+TTGINMNYLLV-CRNWYAMCLP--KLYYAPA LRR
ScYor080w=Dia2    L= 732  207 VGNLPIEILPIIFQRF-TTKELVTLSLV-CNKWRDKILY-HLDCFQEF TPR/LRR
ScYbr158w=Amn1    L= 549  166 Avfeipeiveniikmi\kqgalfscmmv-nrlwlnvtrp---flfksl LRR
ScYmr094w=Ctf13   L= 478    9 PSFNP-VRFLEL--PIDIRKEVYFHLD@-YAFWLRYDCLV-------- LRR
ScYlr352w         L= 807   15 gaaippeivyqiltyq|lttfvksnltv-nktfshicqv---liyryc
ScYbr203w=Cos111  L= 924  187 INDLPVEIIAKILSEF?DQKTLVRCLYV-SKKFYKATKI---VLYRLP
ScYlr224w         L= 369    8 LMDLPLEIHLSLLEYV--PNELRAVNKY$SLAWIAEDN----YIWAVV
ScYml088w=Ufo1    L= 668    8 LQDLPPEILINIFSHL-DEKDLFTLQEL-STHFRNLIHD--EELWKNL
ScYjl149w=Das1    L= 663   49 LTKLPDELMQEVFSHL-PQPDRLQLCLV-NKRLNKIATK---LLYRRI
ScYdr131c         L= 556    2 FDKLPYEIFKQIAWRI-PQEDKISLTYV-CKRSYESIIP---FIYQNL
ScYlr368w=Mdm30   L= 598   16 IDHLPPEIWLCISKLV-GTSDLHNLCLI-NRRLYLTITS--DEIWKRR
ScYbr280c=SAF1    L= 637   12 DAGLSPDIVQATLPFL-SSDDIKNLSQT-NKYYNTLLDFD/KILWHEL
ScYdr219c=Mfb1    L= 465   17 LTNLPLNLLFRILSHL-DMNDLQNIGKT-CTLLRMLANE--NIVYRNA
ScYnl311c=Skp2    L= 763   57 LMCLPTKVLLLILRTL-DFNTLVTLCQV-NSRFYNLITN--EFLFQNV
ScYmr258c         L= 553    1 MAFQDQDIFIVFSHAS$NQNDLLSLSLT-SKKMHDMIAI--PRLYSNI
ScYlr097c=Hrt3    L= 344  100 LEILPDDILLRIIKKV^SGESWVNLSMT-CSTFSKLCFH-DSVPFKTF
ScYjl204c=Rcy1    L= 840    3 DLLKVPEIVTNIASYL-STVDYLSFQQV-NKRVYAIINGK/SKYWSLK
ScYnl230c         L= 379   22 VSNVPYHLLKRILQKV-KIPQLLKLEK--SNVLLIFDDD---ELWLEF
```

^#^ Acidic residues are in blue, basic in dark red, small in red, and hydrophobic in green. Positions matching the consensus motif (panel A) are highlighted in yellow (hydrophobic), gray (acidic), green (basic), or teal (small). Bolded residues correspond to the PxE motif.

Residues skipped for the alignment are as follows: /=HS; &=LFL; ^=ILM; +=DIPEKEEKLNKTANGKKT; ?=ELGR; $=FYVLHNHSYKEK; @=76 aa deleted; \=57 aa deleted; |=18 aa deleted

**C. Candidate F-box motif sequences from *Toxoplasma gondii*** (type 1 strain ME49)

```
Name	 Gene ID*    Length Start  Sequence‡					        domains
                      a.a.
FbxW1  TGME49_261370  1618  941  WTSVPAEVLSTLCQFL-AVEDLVAFQRL-DRRAYAVGSH--ATVWRAL WD40
FbxW2  TGME49_299230+  636   16  PACLPTAIVFHVASFL-TPNDVCSMGAT-CRKWREVCCSDLQPLWREF WD40
FbxW3  TGME49_110910^ 3679  236  LESLSPCLLANLVSFL-LPLDIVSVSLC-SRSLLWLSRC--PFVWRRC WD40
FbxL1  TGME49_262530   979  173  FAELPQEVLELIFSRL-GLADLSRCLCV-AKSWHPPLNA---VFAKTI LRR
FbxL2  TGME49_313200   839  307  LSDLPEELLQQILDCC-PKECLLVSHAL/RRRVLRLTPL---QCTEPR LRR
FbxO1  TGME49_310930•  808  444  FPFLDEPALSLLVPFL-FGRSLATCMTV-CPHWFMKINR---AMERMC
FbxO2  TGME49_215210+  683  254  FNTCPAECLQAVFHFL-HVEDILRMQVV-SSAFFSTIRDEIGAFTHIR
FbxO3  TGME49_225900+ 1461  160  LEALPAGCMYTLFAFF-DVSEVAELRLL-SRTVKAVVDS--PCSLRGC
FbxO4  TGME49_228380¶ 1726   42  LCFDYPLCFDYLLSFL-DLRDFLTLSLV-SHSLRDILLSDLTRAARCV
FbxO5  TGME49_243750+  865  478  LCSLPQELLDVLPLYL-DAFALTRLSSC-CRLLHRLCGNRSDVCWRAK TPR
FbxO6  TGME49_258900¶  577    3  DLLQHPDIVGRILSCL-AWRERRQLAAV-CVSWREAAET--SPCWSDL
FbxO7  TGME49_275780   978  211  IDALPDDLLCEMLLFL-PFDEVGASIPLVSRRFCRLALL--PYIWTFF
FbxO8  TGME49_305630  2045   61  SPDGPCETIVFLLGFFLPVADLCSCSAV-CKAWWAVCTLQHQQLWRER
FbxO9  TGME49_359350¶ 1788  742  LRQLPPAILCAVLRFL-PCASVLAFGTT-CRYAHDLVQL--PAAWNLL
FbxO10 TGME49_215620¶ 1930  280  GAEAPEDARGCLLAFL-TWNDLARLRAV-SRQLKRLVEN-AALSARAV
FbxO11 TGME49_203040^ 1894  311  FSDLPDVCVRICFSFL-SVEEILKYQFL-SRYIRRAIGL---DHVLPL
FbxO12 TGME49_278815¶ 2186   84  LHLLPSSALSLLLRFL-HLDDVCRLALS-SKQLYLHPDLNTPFAVAHL
```

F-box motif consensus sequence^¢^ #xxL***P****x****E***ILxxILs#L-xxxDLLxLsx#-CbxWxxLsxx--xxIwbxl

*Sequence identifiers are from [www.toxoDB.org/](http://www.toxoDB.org/).

^‡^ Acidic residues are in blue, basic in dark red, small in red, and hydrophobic in green. Positions matching the consensus motif are highlighted in yellow (hydrophobic), gray (acidic), green (basic), or teal (small). Bolded residues correspond to the PxE motif.

+= reported also by ref. J.

^¶^= Identified by hidden markov model

• supported by interactome studies

/ = AAAVRR

^¢^ Consensus: #=hydrophobic side chain, s=small, b=basic, PxE motif is bold italicized.

**D. Predicted F-box motif sequences from *D. discoideum*.**

gene	C-term.

name	dictyBase.org ID	GenBank ID	Length	Start	Sequence	domain

*WD40 and LRR types*

```
fbxA	DDB_G0276887 XP_642882.1  L=1247  634 FDNLPEEVVQIIFSNL-SAINIVNLSLV-CKRFKMATDS--PILWKNL WD40+
	DDB_G0285445 XP_638057.1  L=1178  720 FDLLPYEMIQYIFTLM-DATHLIRMSRT-CKYFNRICLD--DNIWRDL WD40
	DDB_G0286129 XP_637855.1  L= 739  294 TLILPEEVMLHILKFL-SPIDLCRGVCRVSLKWRSLAFD--ISLWRDL WD40
	DDB_G0280895 XP_640968.2  L=1501 1029 LFYLPHEIIMTILSYL-DHRALCKVSRT-CKYMKQMSES--DILWYRL WD40
	DDB_G0292834 XP_629376.1  L= 569  137 ISILPIEITMKIISYL-PFQDVLSIQYV-CSEWFLITCE--EILWKSI WD40
	DDB_G0290127 XP_635883.1  L= 649  132 FDNLPSEVIEKIFSYL-FEYDIYRVSLV-CNYWNQIAKS--NDIWLNI WD40
	DDB_G0291716 XP_629997.1  L= 773  294 FEELPLEIIVYILGFL-ECEDIGLRTSLLNKNFRDISDF--YFIWKNL WD40
	DDB_G0280581 XP_641061.1  L= 689  124 LFLLPNEIISNILSIL-EVKDIVALSIV-NNHFYYLINS-YNNLWKNL WD40
	DDB_G0269442 XP_645974.1  L= 923  423 FQLLPLEVMVMIFENL-SFHDVHYCVSLVCRQWHSYTFA--DSIWKNL WD40
fbxD	DDB_G0292312 XP_629697.1  L= 530  114 ILNLPSTIYAQIFSEF-PVKEILKFSLV-CKEFNKAINH--KFLWKIK WD40+
mkkA	DDB_G0283265 XP_639165.2  L=1267  858 NIILPINLIILIFREI-KPNFVNTLSRV-CKHWKQIIDD--DELWNKY WD40+
	DDB_G0281237 XP_640903.1  L=1038  238 IYGLPMEMLFEILSNL-SVVDLVKVSKV-SKFFYSVVNN$TSISFENM LRR *
	DDB_G0268536 XP_647378.1  L= 676    9 ILKLPLELLEKIFKFL-EQKSLCSTSLV-NKQWNHTVNL----VWHTI LRR
	DDB_G0275117 XP_643758.1  L= 638   37 FNLLSTELILYILKYM-GTHDLCILARI-SKRFRECCYD--STLWRTL LRR
	DDB_G0289577 XP_636156    L= 729   98 IRDIPYDLLGKLLali\LTKEFMSTSLV-CRKWRALARL-NGNHFNLE LRR
fbxC	DDB_G0285563 XP_638127.1  L= 283   33 INFLPGTVLLKIFSMV-NENDKPNELP--SQSFYQLSLV--CKLWRYY LRR
dlrA	DDB_G0290485 XP_642416.1  L= 818  282 IGSLADIIISNILGKA&NSQNFTNICSV-CSKWRKISVG-RLVNYTYQ LRR +*

ankyrin, kelch and armadillo types
	DDB_G0290983 XP_635434.1  L=1335 1048 INNLPQEIIQNILNQL-NDIRDLNNCSQVSSSFYRVCSD--QIFWKTK ankyrin
	DDB_G0272020 XP_645384.1  L= 803  138 INDFPEEILIHILKWV-SPFDLRICSLV-SNYWSIISSQ--NPIWYEK ankyrin
	DDB_G0276013 XP_643383.1  L= 748  195 FFDLPIEIRMHILSFS-DAVDLSKTCTV-SKYWKSMVDD--EQLWNNL ankyrin
	DDB_G0285411 XP_638276.1  L=1438    1 MEDLPTETLILIFKNL-KINELLNVGLV-SRFFFLVSSD--DRLWKLL ankyrin
	DDB_G0292070 XP_629810.1  L= 827  287 IHELPEEIIVKIISYF-SLDTIFNMSLV-SMNFYRLTQD--YDLWKKK kelch
	DDB_G0272386 XP_645182.1  L= 447   52 ILEFPDEVLLTILSFL-EPNDIKNIYLS-SRYLSSFCFE--NKIWKAI kelch
	DDB_G0268696 XP_646870.1  L= 543  103 LPNFPEEIKLQIFSHL-SASDLVSISLT-CKTYYAIANE--RTLWRNL kelch
	DDB_G0282667 XP_640191.1  L= 526   34 LNIFSYEVLVHIFSFL-DLTDIKNLKLV-SKDFSNIASE--CCLWRMF kelch
aarA	DDB_G0288877 XP_636508.1  L= 757  314 IFLIPTEMLVHLLSFL-SANDLWRISLT-CKRIWYIVDV--FKFWELL arm

other known domains
	DDB_G0282285 XP_001733003 L= 525   14 ILNIPFENLLIILQNL-HTKSLLELSQV-NLYFNQIISN#SNMIWNDI
	DDB_G0278671 XP_642325.1  L= 534  286 AIIQSEKLQFKCKNRI-NVQSLIQVSST-NSLLQLISND--NELWKKI
docC	DDB_G0292004 XP_629761.1  L=1736  423 ISVLNGEVLVCIFQYL-DIKEICCLAQT-CSWWRIVSEQ--NVLWLKK
gacFF	DDB_G0275085 XP_643800.1  L= 885  351 FSLLPTHLTLYVFSYL-EPKELLILAQV-SSQWQKLAGD--NLLWVRF
	DDB_G0292302 XP_629688.1  L= 933  308 SNGLSDEYWILIFHFL-DFKTLVALSET-CHLFYRLSND--RFLWESL
fbxE	DDB_G0281783 XP_640639.1  L= 781  325 LSTIPEVQLFDILVLL-KPIDLGRLRVT-SKYFATIVAK--DSLWRRI +
	DDB_G0276383 XP_643197.1  L=2026 1637 VFLDSLQVSILFGYLL-DSDSLVACSMV-CKLWRKCSLN--SNIWTRF
gefG	DDB_G0272032 XP_645356.1  L=1556  310 FLDLDEKIYLKIFGYL-FAEDLCSINRV-SKHLCNIINN--QQLWKDL
gefG	DDB_G0272032 XP_645356.1  L=1556  689 SRVSDILLIKIFRNLD-SIKDLSVCQRV-SKRWNKAIAI--SSLWEQL
	DDB_G0288147 XP_636866.1  L=1347  876 RLFNSSIIQLKILKYL-DTNTLLNCGLA-SRQLRYNLYN^FSNFWIKL *
jcdH	DDB_G0280485 XP_641202.1  L= 536   57 SVFEDNFIIDSILDLL-SAEELTKTMNLVSKTFYIYVQE--EEQWKMR *
jcdI	DDB_G0270006 XP_646466.1  L= 983   64 SILEDQILLNVVFNEF-TCSELLKYQCV-SPAFYILLGD--DRLWKDA +*
	DDB_G0278443 XP_642368.1  L= 541  157 LEVLKPELVNYIFLKV-GALDMTSLIGV-NKQCYRIGMN--EKLWLYF *

other unknown domains
	DDB_G0275777 XP_643474.1  L= 843  141 ISDLPYEVLCKILSFL-DVNTLISTSLV-SNLFNKISNS--NEIWKPK
	DDB_G0280141 XP_641265.1  L= 616  230 VNGLPEEVLVRIFKIL-NVEDLYKCYSV-CSLWKRLCED--EAIFRKV
	DDB_G0269446 XP_645976.1  L= 358    3 LQDLPTEVFYLILSKV-SSDFIIQEFQYVSKYIYHLCSD--ELFWKLL
	DDB_G0293826 XP_628954.1  L=1261  505 IGSVPREVWIHILSFL-NERDLLNISMV-DSFLNEISQD--NTLWKPL
	DDB_G0290985 XP_635436.1  L= 789  241 IEDLPFEVLIHIFSFL-KYSDILIVSLV-CKEWNSHSLV-ATTELTIE
	DDB_G0268822 XP_646992.1  L=1032  253 EIYLPIDVLDLIFSQL-SFIDRCICASV-CSSWRNVLIMQPHPLLENC
	DDB_G0289099 XP_636405.1  L= 784   84 ILELPDDILLMIFRYL-DVPTLLKVSQL-CNFLNSVSSN--HELWKEL
	DDB_G0291902 XP_629902.1  L= 336   23 CEELPYDILLYILSYL-TPIELCCLSRV-CYSFRVLAED--DWIWRGF
rliC	DDB_G0276149 XP_643304.1  L= 328  172 SNILSKEIQSLIVLNL-SIYDLLSLFRV-NKYWYEFGQN--NSFWKSL
	DDB_G0285125 XP_639880.1  L= 540  159 LISLPHQILYQIFESI-SPQDFTNCSKL-NKKWNSVLIK-LDKCWIQY
	DDB_G0278767 XP_641950.1  L= 636    1   MKSIVLVFNILKHL-EIDDIYCCEIV-CKDWRKVANQ--NKLWEIL
	DDB_G0270452 XP_646815.1  L= 830   22 KISNNLIIVKEIVSNL-DVLDLVGIQYV-CKQWNSIATS--NVIWLEA
	DDB_G0285933 XP_638013.1  L= 906   60 NDLFSVELMVKIFGMC--DPFSTALILV-CKHWYSVLINNQLSLWRDL *

weak candidates
	DDB_G0287541 XP_637180.1  L= 669  153 EEESISANYNSYNKKT-ILNVVINFSMT-CKYFYYLFQD-DGFWYRTY WD40
	DDB_G0270750 XP_646595.1  L= 485  426 FNHFPEEILYQIFSRL-SQAGKLIPLQL-DYLIRKSYLADQVALVLST kelch
rabGAP	DDB_G0280955 XP_640996.1  L= 733   78 TLLLDSEILIRIFSKL-EYSDLFIVKRV-CSRWKPLCEH--KSLFNSL other
```

Acidic residues are in blue, basic in dark red, small in red, and hydrophobic in green. Positions matching the consensus motif (Fig. S1A) are highlighted in yellow (hydrophobic), gray (acidic), green (basic), or teal (small). Bolded residues correspond to the PxE motif.

+=verified Skp1 interactors

*=F-box motif not recognized in NCBI Conserved Domain or Uniprot databases.

Deleted sequences: \=hgNRI; &=HKDEP; #=SSI; ^=GISMEKKT; $=LPSC
