## Supplemental Figure 2 for "*Toxoplasma* F-Box Protein 1 is Required for Daughter Cell Scaffold Function During Parasite Replication" (from another version of this preprint: version 1, 2019-01-03)

**Figure S2. Homology of alveolate sequences related to *T. gondii* FbxO1**

**Code	Accession	Species	Clade**

[Tg	XP_002364337.1](https://www.ncbi.nlm.nih.gov/protein/XP_002364337.1?report=genbank&log$=prottop&blast_rank=1&RID=0BN63AAB211)	TGME49_310930 [Toxoplasma gondii ME49]	Coccidian A	Apicomplexa

[Hh	XP_008885908.1](https://www.ncbi.nlm.nih.gov/protein/XP_008885908.1?report=genbank&log$=prottop&blast_rank=2&RID=0BN63AAB211)	HHA_310930 [Hammondia hammondi]	Coccidian A	Apicomplexa

[Nc	XP_003885097.1](https://www.ncbi.nlm.nih.gov/protein/XP_003885097.1?report=genbank&log$=prottop&blast_rank=3&RID=0BN63AAB211)	[Neospora caninum Liverpool]	Coccidian A	Apicomplexa

[Beb	PFH34091.1](https://www.ncbi.nlm.nih.gov/protein/PFH34091.1?report=genbank&log$=prottop&blast_rank=4&RID=0BN63AAB211)	hypothetical protein BESB_072430 [Besnoitia besnoiti]	Coccidian A	Apicomplexa

[Cs	PHJ16413.1](https://www.ncbi.nlm.nih.gov/protein/PHJ16413.1?report=genbank&log$=prottop&blast_rank=5&RID=0BN63AAB211)	hypothetical protein CSUI_009775 [Cystoisospora suis]	Coccidian A	Apicomplexa

[Em	XP_013337306.1](https://www.ncbi.nlm.nih.gov/protein/XP_013337306.1?report=genbank&log$=prottop&blast_rank=6&RID=0BN63AAB211)	oxidoreductase, putative [Eimeria maxima]	Coccidian B	Apicomplexa

[Cc	XP_026193228.1](https://www.ncbi.nlm.nih.gov/protein/XP_026193228.1?report=genbank&log$=prottop&blast_rank=7&RID=0BN63AAB211)	LOC34623623 [Cyclospora cayetanensis]	Coccidian B	Apicomplexa

[To	XP_009689732.1](https://www.ncbi.nlm.nih.gov/protein/XP_009689732.1?report=genbank&log$=prottop&blast_rank=11&RID=0BN63AAB211)	TOT_010000887 [Theileria orientalis strain ShIntoKu]	Piroplasmid	Apicomplexa

[Bab	XP_012766453.1](https://www.ncbi.nlm.nih.gov/protein/XP_012766453.1?report=genbank&log$=prottop&blast_rank=13&RID=0BN63AAB211)	hypothetical protein, conserved [Babesia bigemina]	Piroplasmid	Apicomplexa

[Pr	CRH00939.1](https://www.ncbi.nlm.nih.gov/protein/CRH00939.1?report=genbank&log$=prottop&blast_rank=15&RID=0BN63AAB211)	conserved plasmodium protein, unknown function [Plasmodium relictum]	Haemosporidian	Apicomplexa

[Cm	XP_002140319.1](https://www.ncbi.nlm.nih.gov/protein/XP_002140319.1?report=genbank&log$=prottop&blast_rank=9&RID=0BN63AAB211)	hypothetical protein [Cryptosporidium muris RN66]	Conoid	Apicomplexa

[Vb	CEM03254.1](https://www.ncbi.nlm.nih.gov/protein/CEM03254.1?report=genbank&log$=prottop&blast_rank=10&RID=0BN63AAB211)	unnamed protein product [Vitrella brassicaformis CCMP3155]	Chromerid

[Im	XP_004034674.1](https://www.ncbi.nlm.nih.gov/protein/XP_004034674.1?report=genbank&log$=prottop&blast_rank=17&RID=0BN63AAB211)	hypothetical protein IMG5_116140 [Ichthyophthirius multifiliis]	Ciliophora

[Pt	XP_001454827.1](https://www.ncbi.nlm.nih.gov/protein/XP_001454827.1?report=genbank&log$=prottop&blast_rank=19&RID=0BN63AAB211)	hypothetical protein [Paramecium tetraurelia strain D4-2]	Ciliophora

[Tt	XP_001028005.3](https://www.ncbi.nlm.nih.gov/protein/XP_001028005.3?report=genbank&log$=prottop&blast_rank=12&RID=0BN63AAB211)	TTHERM_00498170 [Tetrahymena thermophila SB210]	free living ciliate

[Sc	OMJ83137.1](https://www.ncbi.nlm.nih.gov/protein/OMJ83137.1?report=genbank&log$=prottop&blast_rank=14&RID=0BN63AAB211)	hypothetical protein Stecoe_15990 [Stentor coeruleus]	heterotrophic ciliate (heterotrich)

[Pp	KRX05854.1](https://www.ncbi.nlm.nih.gov/protein/KRX05854.1?report=genbank&log$=prottop&blast_rank=16&RID=0BN63AAB211)	hypothetical protein Ppersa_03791 [Pseudocohnilembus persalinus]	pathogenic ciliate

[Sl	CDW85380.1](https://www.ncbi.nlm.nih.gov/protein/CDW85380.1?report=genbank&log$=prottop&blast_rank=18&RID=0BN63AAB211)	UNKNOWN [Stylonychia lemnae]	Ciliate

[Ot	EJY65153.1](https://www.ncbi.nlm.nih.gov/protein/EJY65153.1?report=genbank&log$=prottop&blast_rank=20&RID=0BN63AAB211)	hypothetical protein OXYTRI_14697 [Oxytricha trifallax]	Ciliate

[Sm	OLQ08676.1](https://www.ncbi.nlm.nih.gov/protein/OLQ08676.1?report=genbank&log$=prottop&blast_rank=8&RID=0BN63AAB211)	hypothetical protein AK812_SMIcGEnE7829 [Symbiodinium microadriaticum]	endosymbiotic dinoflagellate

```
           *
Tg  1     MGNTESTAAELHDNYLAVLKSVEGRLGQYPSRTLSPRSQSSGVDRSRA-RVRPEEAYRDPGKRG-AGNRPRNLPSKGEK  77
Hh  1     MGNTESTAAELHDNYLAVLKSVENRLGQYPSRTLSLRSQSSGVDRSRA-RVGSEEAYRDPGKRD-AGNRPRNVPSKGDK  77
Nc  1     MGNAESTASDLHDNYLAVLKSVENRLGQFPNRSLSFHSQPESNGRSRV-RTNTEEACQELGKRD-AANRPRKLSSKGEK  77
Beb 1     MGNTESTPSDLHDNYLAVLKSIETRLGKSATSESSRRRASGAGGWSRAARKQSEETRPPSGSRERASSRPCKSGSKLDK  79
Cs  1     MGNTESAPPALHDNYLAVIASIEGRLGHLPARNSHAPTKSPR---------------RANGVCH-HSSGPAAAVLPRDG  63
Em  1194  MGNHPSHPSIPLDGYLAVRASIEARF-------------------PPSQELRQIKPTEATACWGAPGGELHATLSGGRP  1253
Cc  1     MGNRPSYSSVPLDGYLAARASIEARF-------------------SHAAASRQ-RTASPSGCCGT-SEDTFPTMSAMEH  58
To        -------------------------------------------------------------------------------
Bab       -------------------------------------------------------------------------------
Pr        -------------------------------------------------------------------------------
Cm        -------------------------------------------------------------------------------
Vb        -------------------------------------------------------------------------------
Im        -------------------------------------------------------------------------------
Pt        -------------------------------------------------------------------------------
Tt        -------------------------------------------------------------------------------
Sc        -------------------------------------------------------------------------------
Pp        -------------------------------------------------------------------------------
Sl        -------------------------------------------------------------------------------
Ot  1     -----------------------------------------------MYSNFSKEQQFQQQQQNNFQQQQFLQQHHTRI  32
Sm  20    VGNLRRQLQQRALRFPEELCVGKTLKLELLQQVGSEVVVCSHEESYRVAFEAGQLALTVSPESGDQDDAASDERFEMDG  103


Tg  78    PSR-----SAH-KSG-STSSVDQFNNG-PGANRKHGKDSVQRRDDAPP-QADQEHSSLQPA----------DGKRRPAAK  138
Hh  78    PSR-----SAH-KSG-STSSVDQLNDG-TGANRKRGKDRVQRRDDAPP-QAAEEHSSLHPA----------DGKRRPAAK  138
Nc  78    PSR-----SVQ-QSG-LNSSVDHSSAS-ADA-KKRGKERAHRRRDAA--QSDEDGNSSHPS----------DGKRR-TAK  135
Beb 80    SAS-----ALSSKSF-ADGNQRCSSES-LGS-KKRGKELAHRRQDTLP-HSDDEKKSSAKST---EGRRGHDAKRQ-TGS  146
Cs  64    PKC-----PVPDNSE-ISGTQSPGTGG-LDS---TGTSESSPRRNSGT-IVPQERRGKPPLT-------GDPGQVEGLGE  125
Em  1254  QEN-----PVA-GEP-CRSNVPLLLNR------------------------VAAGALVPPTL------------------  1284
Cc  59    HPC-----SPH-REH-AKFSREMCLEQ-RGA--------------------VSSGLVLPPSL------------------  92
To        --------------------------------------------------------------------------------
Bab       --------------------------------------------------------------------------------
Pr        --------------------------------------------------------------------------------
Cm        --------------------------------------------------------------------------------
Vb        --------------------------------------------------------------------------------
Im        --------------------------------------------------------------------------------
Pt        --------------------------------------------------------------------------------
Tt        --------------------------------------------------------------------------------
Sc        --------------------------------------------------------------------------------
Pp        --------------------------------------------------------------------------------
Sl        --------------------------------------------------------------------------------
Ot  33    PSQNNNNSLGAGSMSQSIPNLRSFVEQRLNINQLEEDDLVSRGSETSNLKKPQFQKQGAKTLKTSDQNYKKNLFSAMTSR  112
Sm  104   LPMRRQDAAAGAKEYLEKHELLPYLQGLLETVCKEKPANPYRFLWRQLGLALDTSPPKKPAQNP----TGPTGKAQSARE  179


Tg  139  GETPNVESTATGSDWGDsAVNDF--AEEDSCAVESPAHFKL-qTKESGVKAPSPRLFPCETLYEVTSQIDLaDE  209
Hh  139  GETPNVESTATGSDWGDsAVNDF--AEEDSCAVESPAHFKL-qTKESGVKAPSPRLFPCETLYEVTSQIDLaDE  209
Nc  136  GEPPNLESTATGSDWEDaAVNDF--AEEDDCAVASPAHFKL-qTKDSGVKAPSPRLFPCETLSDGPSQRDLEDD  206
Beb 147  SASGVNESTAAGSDWEDsAVNDF--AEENECVVPSPVHFKL-ETKDSGVKGRSPRLFPCESELPSITDL---DD  214
Cs  126  GQGGDRDLAATESDWEDaTVNDF--AEEEVTSLSSSAHFKP-qSKQSSVKAPSPRLFPSTSALKLSSDSD--DG  199
Em  1285 ---SVAETTDTNSDWAPDEFEaA--ASSEILGGPSQSFTRE---KAPRVRAQSPQPFPSMS------------- 1337
Cc  93   ---SVAETTDTNSDGAPDEFEGA-qAAEDMFSRLSQSFTR---VKAPHVRARSPLPFPSMA-------------  146
To  1    ----------------------------------------------MSNYTPTNRRIRCFSFRRRRNVFDYVNV   28
Bab 1    ----------------------------------------------MANYRGFFLRFKGLCARRMDA--SLVRE   26
Pr  1    ----------------------------------MGNKISEcKHRQNILYKNIKNIIKQRNDIDVKNNIYIcES   40
Cm       --------------------------------------------------------------------------
Vb       --------------------------------------------------------------------------
Im       --------------------------------------------------------------------------
Pt       --------------------------------------------------------------------------
Tt       --------------------------------------------------------------------------
Sc       --------------------------------------------------------------------------
Pp       --------------------------------------------------------------------------
Sl  1    MNTNPSIELFNNLDYQRPsKLINEMnDCLQKLDLDDSHNLEP-QSFEDLNGTIPCDFIPNSMTMSQACL--qST   71
Ot  116  QQQESQLNNSQLMMSQRsLtSLHQHDqSKLNQSQIYGGGQLNGLSANKNESQFLHLNTSQNINKSQFLQRsqSP  189
Sm  180  PEKSTRPGTPAELEGsRaTTPLPPPLRPTSEEAGTGCVVSA-DVLPEPLKSEIQEALEAPSESHLQWLHAVaGV  257
```

```
Tg  210   KTQVED[1]ELSSLPSSP     TAS   AEVATDKPGEGR[9]GRSAEGDRGP[9]AD   AKEN-ESDNSHGESDQDL  287
Hh  210   KTHVED[1]ALSSVPSSP     TAS   AEVATEKPGEGR[9]GRSAEGGREP[9]AD   AKEN-ESENASGESNQNL  287
Nc  207   EKTEVD[1]SLGSSPGSP     TAS   AGSVTGATG---   GRSSDARKER   AE   GKGK-DPVNTSDKSKRDP  263
Beb 215   EETQAG[3]ARESAPETP     SAF   AEGAEAQER---   ARKNSVSKKE   AK[5]SRGD-KHARHSGEQKAES  278
Cs  200   RGEEAD[1]AFHTRDDER     NGM   ATEEPEKEKE--[6]AAQIDSSLSS[9]GN[5]VTGD-SSESKPNPDMVAK  277
Em  1338  ------   -----HCSS     TVS   RKAGTHKQQLPE   -REISGSSSR   --[1]SRSN-SSGSNDGGSACST 1383
Cc  147   ------   -----NCSP     TVS   KGSDAQQAPLCN   -SSIAIINER   --[1]YTLN-SERSPNAVGC---  189
To  29    PTQATL[2]LHQPTNTQR     E--   EPPEPSKHDDNR[5]----------[5]EP[7]-----YHPYVTNYHSTEK  90
Bab 27    PTVAVP[2]--EPRPVEQ     T--   EVVEPSTTGDAS[9]PIENAQSPSS[9]EE[9]VTEDVYDPLIMNFSPKIS  111
Pr  41    NEKDLF[5]IFELYNTHE     ENT[1]KKDNENKENEEN[9]KNEKEKKKNA[9]SE[9]KRDDEGKNVKEIDEQTKK  133
Cm  1     ------   ---------     ---   ------------   ----------   --   ------MGSVTSAPSHDK   12
Vb        ------   ---------     ---   ------------   ----------   --   ------------------
Im        ------   ---------     ---   ------------   ----------   --   ------------------
Pt        ------   ---------     ---   ------------   ----------   --   ------------------
Tt        ------   ---------     ---   ------------   ----------   --   ------------------
Sc        ------   ---------     ---   ------------   ----------   --   ------------------
Pp        ------   ---------     ---   ------------   ----------   --   ------------------
Sl  72    PHLFR-[4]GLECTQSSP     QIN[4]SQQIKAALFGGT[9]KIQQQNKDDD[9]LK[9]QVQSqFSTNYKQKNPYKS  165
Ot  190   NNIYQP[5]GNYASQFDP     DLF[4]AMKKQYSQGVAQ[9]YYIQQNDDKI[9]QM[9]PLQEVIPQNQNQNGYAFK  285
Sm  258   EDFENV[5]ALFQSSIQQ[151]TAS[3]KVSAQEPPELAR[9]LTERILCRTA[9]EP[9]REQDaEDQGRGGKEDMPE  503


Tg  288   RRRSDGRGASSQRSR--------DEASNSGEPPSQSKPSQL   SKQIRKKELAPEPLRAADIDLSSDVRVGASLKGGK    355
Hh  288   RRRSDARRASSRGSR--------DEASNSGGRPSQAKPSQL   SKQIRKKELAPEPLRAADIDLSSDGG-GASLKGGK    354
Nc  264   RQRSNGSVASS-GTS---------GACLHTGQPHAARKSSP   PSKQSRQKEQVTQLSAADIALPSHAS-GGCTQEGK    328
Beb 279   LEREQASGSER-GSP---QKKPsRGLSCSEARRSSAKNSAS   LTNGRKKERQPQALCAEEVGLSSAAG-TGVRKKGS    349
Cs  278   LSETKVGRSLS------SKKStnSLAGTSTTRPSKSPAPGQ   PRSRGPGRNVPEPLRAEELEYAVTTA-SVCSRRG[11] 356
Em  1384  RKVYAVSAFPGPSSR----PKQLDATLVASSGGAQRPRQDR   SQQKNSSRSPPGACIADQHKAGFPVG---TRQTR[ 3]1454
Cc  190   ---------PQPKRK----PQQPAEALIAPTAG--------   -----------------------PHQTKQLGTEA     223
To  91    YAKKPTSKKVTPKSSIGEPPPPPKPPLNHAPSGPKYLE---[4]DLQTDFQKLSTETITR------------------     148
Bab 112   RLRAEAADSDTCKSPtRDVNSIsQAAATSTSSGHTAEQGVF[4]---TLFSDVGSKPMAE------------------     169
Pr  134   TNENHKKNFELDYGSFYNIEEKnNICKNNNNNKLLKKEKLL   -KHIEVKNEKSQKLKNTEKTENVLLK-KISLNKN[ 9] 215
Cm  13    HVYPKNNNSIEPLSRsYSNVELPSVTPVSRSQSIQTLSAPL[2]DSVSDLIDRISSRLNKNYRIVKYLAL--ACPKDY[ 8]  95
Vb  1     ------------------------------MGNVSSGSSRF   DARKEDRKFKIRCSIIKRLGPPGAPMPETVPPPS[ 7]  52
Im        -----------------------------------------   ----------------------------------
Pt        -----------------------------------------   ----------------------------------
Tt  1     -----------------------------MGIDISKPNQTQ[4]NSNLQF------NNHKSETIQKMILRVQSQIEKG[12]  56
Sc        -----------------------------------------   ----------------------------------
Pp  1     ----------------------------------------M[4]TKQLEYQKEKIYNKDKNDVINKMIQRVQINLEK-[ 4]  42
Sl  166   RQIS---IHTNAYLAtDVG----DSTSHKQEIFSTAQTTQM[4]NNHQRVASNT--KSFFNQTNNNTFISYDNNMINE[ 8] 243
Ot  286   RATNDSVMLSNSFHNtqAPLPKVGQNQYQNQIQDFEDSQAQ[4]QQQLLSNYERAKNLDLSQ--QNKTHTLNSSGKK-[ 6] 367
Sm  504   AASQGQGGTSPEARAKRLMELRRSFSALKQSHGLLQRKVSK[5]DEVRRNTDQAVQLLRDAFVSQRSESD--SKEEKG[10] 591
```

```
Tg  356   GGKGDAEGDNVADLTAGFEGFSPYSPSYSLGFSQPSSKK--AAKTTAENSKAHLPG-GTRGSGNNTAVD---RDACIYGN  429
Hh  355   GVEGDAEGDTVADLAAGFEGFSPYSPSYSLGFSQPTSKK--AGKTTAESSEARLAG-GARGSGNNTAVD---RDSCIYGN  428
Nc  329   E-KGWADGD-LAEFAGDFEGFSPHSPSYSFGLTKLPKKA--IAKSGKTEERPPSAGTSARGSGNKASAD---ADACIYGN  401
Beb 350   K-ELTAEKG-GSAAAAQFEGFSPYSPSYAFDAPTPPIKA---TTTGRTEARASCRG-GGRGDGGSAGPEED-ADSYIYGQ  422
Cs  357   GASSTQEKR--PVSAVDLDSFSPYSPSYALGITTPTTNA--TTTTGSTEDRGSSRG-GFETEGDRLGS-----DRLIFGD  426
Em  1455  GAPEGRAAGVPSAGAEEHEMFCPLSPNSALGSCTPSK----ASSKSAPKTMGQTAV-FVGTTSSSHGMPA--ERNISYGP 1527
Cc  224   GTPSGRPSGLPKACINESDMLCPLSPNSALGSISPPAVAFKGASAKSTETLPETGA-PVQGTSKRF--------SVSYGP  291
To  149   ---SSRKGSVTY-----------------HSFTREDHAP--SRDVLGVGSRFLIGGKLSDGKPSMESRVWDMIERRGWG-  205
Bab 170   ---SAQKNHVS------------VLDDRIKTRTSPDFSP--S--------------KLAKCRSSL--RYYDLMGHR----  212
Pr  216   DDESTFFNDENNNTILTKNSHMDNKELFKSTFKCIELKLLPDDDSYTGNIKSNIAEINYSNNIKIKENKNEINENLKNNK  295
Cm  96    EENTYVKKDEKFKWLNLFSHNLRYSGYDLQTGDKNSEILDtGNETGALNPSSKQHCLGSTCQFKGHSLYEDSKLYVVYGH  175
Vb  53    NLFGSAV--------------------SAAGASTPTS----------------SSGRGRSHSPRSSLPPPPPSPSAYYGH   96
Im        --------------------------------------------------------------------------------
Pt  1     --------------------------------------------------------MQRKESKKATIPKMMARVLQQQEA  24
Tt  57    SMQGRQQIQQQQQQTQNQYYQTPQGQMQMQQQQRFQNMQYQPGNYQQVNGRIIQMNGQNQQPQYGYQNYQQLKIKVNCNV  137
Sc  1     -----------------------------------MGITNTKPKDSEEVITSREYRSELIQDQRKRRHKTFAcMVQRVLI  45
Pp  1                                                                                  MTK   3
Sl  244   -STNPQNQNNQIKQQIQQEKPRKKDKKPNNDDEFVNNYK----LALQQYQEDMNKQAYSHKKRLVSYDQDLKLRQNLRRK  318
Ot  368   ESHQILPSNTVPDSSQN--QNSNLSNVKERDSMLPPYKI-----KIKGNSNMKQEQLLQLTNRKSSSNLYTNYQHPIQRV  440
Sm  592   RIRKHPASGASLVEAMGGTSSQPIDVRVTDRKSLTKELCLRRKDYESMQHRYCVEDLTNSQENSRSRMAHLLHRAAIRDP  671

                   .           Predicted F-box domain        .
Tg  430   -----KKNHFPFLDEPALSLLVPFLFGRSLATCMTVCPHWFMKINRAMERMCGP-ATKGFQQMYSKYLEVWGSA--  497
Hh  429   -----KKNHFPFLDEPALSLLVPFLFGRILATCMAVCPHWFMKINRAMERMCAP-ATKGFQQMYSKYLQVWGSA--  496
Nc  402   -----KKNHFPFLDEPALSLLVPFLFGKSLATCMAVCPHWFMKVNRAMEHKCAP-VTKGFQQMYSKYLKVWGSA--  469
Beb 423   -----ERNHFPFLDEPALSLLVPFLFGKSLAACMAVCPHWFMRINRAMERMCAD-VTDGFAKMYANYLKLWGGA--  490
Cs  427   -----EGNHFPFLEEPALSLWVTFLLGKSLAACMAVCPHWFMKIDKAVSRMCAP-ITRDFAQAYSSHLEVWGSA--  494
Em  1528  -----ACNHLPFLEGRALQALLPFLFGRTLGICMSVCVHWFMRISNGLAEMCSP-IVRGFEASYGKYLYPVNAT-- 1595
Cc  292   -----LCNHSPFMEGRAVQILLQLLLGPSLGACMGVCVHWFMKISDALADMCAH-LSKDLEASYGKFLSPVSAT--  359
To  206   ----MVHENWNMVGARAFNSIVKYLTGMEISLCRQVCKGWYQNVTDVMVMRSEV-IVSAFKKAYKEKLEYDRSY--  274
Bab 213   -----ISEDWADLNSRVYDALLQYFTGSDLTAGQKISKQWSDNVRLLMGKKTAK-IVAAFKVAYQGQLDFESSY--  280
Pr  296   ---ILnKFLNLLKEKKILQNILSFLNCRDLLSFQKTCSIIYIYVSDFLDYICLN-IYSNFKRIYDFYFTPFNFF--  365
Cm  176   -----LNNHKPFLSPGIIDTFLPFILDETLLTCFTVCPHWFLTLVCHMNQVYSHLIDDQFKENYKGYLELEHAT--  244
Vb  97    -----RANHMPFLEPAVMRRLIPFLWGQSFVRCTAVCPQWYMMLLAGVDDILQE-VDGKFDRRFAPYLKMEHTR--  164
Im  1     ----------MKLPFLAQNKIFFFLF-DEQINIKLINSLFYYKFSEVLEFEMVN-LDNKFIKNTMSFLSFKDS---   61
Tt  138    NCNVNEYLFQKLPSSVLYMILSFLM-NDYISIIMVSPLWYYKLHECIETELLK-IDNDFIKSYSNALLFKNS---  203
Pt  25    TDVLTRFLFIKQGQDRIRRNILGFLI-ADFTQLILVSGQWYVGFHQTLKEWLED-LDNRFIKAHLHILSFKNSDFL   98
Sc  46       TPSKTLIDTIPNDIFFIILQFLT-PEIPTLLAVSPKLHVKVYETIDTAFNA-IETQFAIIHSNLLCFKKSF--  114
Pp  4     QLEYQKEKIYNKDKNDVINKMIQRVQ-INLEKNKENTRPSNNAVNNQMESFRVPSMDSVFNNDNTNLQNKMTT[66] 149
Sl  319   --IKSVVQDPAQVPMNIFYTIISYDG-KLFKKFLCLSNGMKNHILFYVFDKTKK-ISHDFHRKYGQLLELQSRS--  388
Ot  441   ---IQFGGNNCPIDTEIVMMILLFDM-PNFRKYLCLTPQWHHSILEAMDEYFKK-VECNFVMKNYEYLMFKKS---  508
Sm  672   EIcKGGENSQIFMEAGMLPLVISFLFGGALIRCFEVCPLWFVVYYKTLDQVFSK-IDKGFKQYYGQSLHLEYAR--  744
```

```
Tg  498   -VKLQPLQTVGDGGVRVDWVIFAKVLPECE--GHILDISYTYSYIS[43]LSPCGQP  590
Hh  497   -VKLQPLQTAGDGGVRVDWVIFAKVLPECE--GHILDISYTYSYIS[43]LSPCGQP  589
Nc  470   -VKLQPLQTAGDGGVRVDWVIFAKVLPACE--GNILDISYTYSYIP[42]LSPCGQP  561
Beb 491   -VKLQPLQTADDDGVRVDWVIFAKVLPRCA--GNILDIGYTYSYIP[44]LSPNGQP  584
Cs  495   -VKLQPLQISEDDGVRVDWIIFAKVLPRCA--GNVLEVGYTYSYLP[42]LSPSGPR  586
Em  1596  -VKLQALHTTEGNGVRLDWVITAKVLPPSA--NHVLSLGYCFKYRP[50]RPPSCAL 1695
Cc  360   -LKFLPLQIAGAKGTRLDWVITARVLPPAA--DNVLSLGYCFEYR-[47]LRPFSRT  455
To  275   -ILFQPVLTA-TKSLRIDVIIRAKIQPYCT--LHKNSFGYTYSYQS[49]-------  365
Bab 281   -MICQSVFTA-QKGMRIDLIIRARVLAWSV--NSKNTLQYTYSY--[ 5]-------  330
Pr  366   -YKYEYLYTD-KPSFRLDCILISKIKKGCI--GYNNRFGYKYNYIY    -------  409
Cm  245   -VTMQPVLTV-CGSVRIDRVLYAKVLKSCA--KKCVTIAYNFKYDK[29]-------  319
Vb  165   -VEWTQVHVA-DPGIRIDRIIVARVTEKCV--GRTITVGESFRLLP[111]------  330
Im  62    YVSYKTQKFNNIKGIRMDRIINAELLPNIK--SNIIYLFYYIFYFL[19]-------  124
Pt  99    QTSFWVDSTKTKKLFRWDRTIISEVMSGFN--GKCITIAFKYNHK-    -------  141
Tt  204   YLTIKPIKVNNHFGFRMDRNICAEVLPSIS--NKTVSIKYTYKSFR[ 6]-------  253
Sc  115   -TDFTRMTVSDKKGIRIDRVIVAEILPYLN--GKTLKIRYNYRQAH[ 4]-------  161
Pp  150   HVSMAPIKISKNKGFRMDRNILAEILPTIK--NKCVTITYTYKIFQ[ 8]-------  201
Sl  389   -LIVKPLSFGKEEGVRLDLSLRVKIKKEAK------KIQGKSIIFS[13]-------  440
Ot  509   YTNSSLIHFCGQKGIRVDRIIVCEVLNSNKMLNKCLRISYSFKYKS[ 4]-------  558
Sm  745   -TDWSPVHVA-EAGIRIDRILIARVTSHHV--NHCTKVSYGYAYKS[13]-------  799


Tg  591   CPTNMPS-RVFTVSYSLAAGAAKSSRTLWMHRDLCRF    HG-DETGVAAMGSVSSVCVGDFVEVAVTVYNGGGR  660
Hh  590   CPTHTPS-RVFTVCYSLAASAANSSRTLWMHRDMCRF    HG-DETGVAAMGSVSSVCVGDFVEVAVTVYNGGGR  659
Nc  562   CPTHTPS-RSFTVSYSLAASAAHSSRTLWMHRDMCRF    HG-DETGVAAMGSVARVCVGDFVEVAVTVYSGGGR  631
Beb 585   CPANSSS-RSFTVSYAFATSPANSSRTLWIHRDMCRF    HG-DETGVAAMGSVARVCVGDFLEVAVSVFSGGGR  654
Cs  587   CPSHTPK-LTFTCSYAFAVSPANGSRTLWIHRDMCRF    HG-DETGVAVMGSIARVCAGDFVEVAVSIYSGGGR  656
Em  1696  GEATEH--RREAPMFSVAVAPAASKRRLWLHRDMCRF    HG-DETGQAVSSHLGPVCVGDFVEVPVVLSNGIGA 1764
Cc  456   GESPKAGCRREVPTFSVAIAPAGSKRRVWLHRDMCRF    HG-DETGQAALAPLGPVCVGDFIELPVVLANGIGI  526
To  366   --GKKSV---FISKFIFEVLKKNSCRTITFTRDLSSM    HA-DDINLATSASVCQVCEGDYIEVPITLISAIGA  431
Bab 331   --ARQPT---YANRFVFECLRRNTRRTVSFMRDVSSV    HG-DDLHVATAESVSQVCEGDSIEIPVVLMNALGN  396
Pr  410   ------RKSSYYAYFNFNVLKKNSSRIIEIHKDISYN    NG-DDINVSHIIN-NYVCSNDYICIPINLYNFIGN  473
Cm  320   --KSSDINEAYLSIYKFEALYTGTRRKQWAHKDISRC    HS-EEVCVAQTTMIPSVNVGDRIEIAVNLSNSFGV  388
Vb  331   GPTSPPRIGPFEYRYKFQTLPAGRSRQLWVHRDMCRF    HG-DETRVATVQNVATVCVGDRVEIAVPLYSPRGC  401
Im  125   ---TRKKDQIYSCKYQFDAIETNKNRIIWAFKEECKV[17]HNYDQQKQTFVQAITPIKVGDNIQIAINYYNLGGM  210
Pt  142   ----------YRSQYKFDVLPQNSKRVIWIAREQTK-    HNFES--VTQVMNMQPILSGDCVKIAINFYNKMTF  200
Tt  254   ---SQQKQKKYLCEYHIDAVQKKSSRTIWAYKDDCN-    HNYDKKKVAFIQPITPVKIGDNIKISVNFFNMNGL  321
Sc  162   ----------QKAEFKLDCA-SQSKNTIWVHRDECKF    HG-EDNKRAFTQQIPLVNIKNNIEIAVNWYNLSGN  221
Pp  202   ----PPQQQKYQCQFKFDVVQ-KKHRYIWVYKEECK-    HNFDDKKYAFVQPIQPVVSGDVLKFSLNFYNLNGL  267
Sl  441   ---ESKIEGEYQNVYQFDLYCARANRWLWLHQDECLF    HGsTDSR-AYSIPIVPICVEDNIEISLTLISLMGV  508
Ot  559   ------SKQTFYADYKLDVVKPNSERVVWLHKDEQEQ[14]---DIMNRPYIQPITQICAKDTIEFAINLYCLQGL  635
Sm  800   ----EDRKDPFECSFKFDTYTRGRSRNFWIHKDICRF    HG-DETRVAATQNITPVCVNDRIEIAVNLFNAMGL  866
```

```
Tg  661   VALDKVKWLPARVEWRREAV---   STRGVFNR---  EICPLERCS---PDWLPADQFRIMTTE   RLKAPEDF-SP  722
Hh  660   VAVDKVAWLPARVEWRRDAV---   STRGVFNR---  EICPLERCS---PDWLPADQFRIMTTE   RLKAPEDF-SP  721
Nc  632   VALDNVHWLPARTEHRQESV---   STRGVFNR---  EICPLERCS---PEWLPADQFRILATE   RPKAPEDF-SP  693
Beb 655   VALDRVCWLPPVQERRRDSA---   PSRGIFSR---  EACPLERGA---PDWLPLDQFRVMTTE   RLKDADDF-SP  716
Cs  657   VALDRVSWLPLRQEKRRQSL---   PSKGLFSP---  EACPLEWST---AEWFPADEFRLMSTE   RVKDPEDF-GP  718
Em  1765  SDVSSIQWFPISIARRVPRV---[9]ASKGLY-----  EGCSLELQY---TEWFDGDQYRHMTTE   RLQRAECL-EP 1833
Cc  527   ADLHTIRWLPMHVAPRLSRV---[9]AALGLY-----  EGCGLEMQY---VEWFDGDQYRHMTTE   RLKRAECL-EP  595
To  432   TDVKSIKFLPIVRDVLSKAP---   SHLDLLHR---  EWYTVEPNS----KYLEQLFPQNIHSY   SIIHPHKL-LP  492
Bab 397   TNVDSIKFLPMRREAVEGEP---[1]SSLDKMHR---  EWYTVGPNS----HYLQQLFPADIVAY   NLVQPEKL-QP  458
Pr  474   VDFNSIKFISNKLSKYIIY----   -----------  NNQLDDQM-----WYNKEEYQILIKEN   RLITFESL-LP  524
Cm  389   VNIKSIEFQPISFETIKSE----   -----------  N-CEVEDLYPNTSDWSSIDNQNEVVES   --FHIPDF-SP  441
Vb  402   VDMQTFRWMPLRGEPATKQI---   -----------  --FPLEKELD---DWYELDLFQSQTVE   RLSVPNHF-EP  453
Im  211   IDLDSITWKEPIFQHIPKIPI--[5]-----------[9]ICEIEQIY---SGWILYNYFHNNQ--   KC-RLDFF-EP  275
Pt  201   IDPDTIEFEEPSIEQSKEC----   -----------   ICPIQSLF---FDWVGIDYAKQKP--   --KLNDYLVHP  249
Tt  322   VNIYSIQWKELTTGSIPSVPI--[5]-----------[9]ICETEEIY---SDWVLYEYFHKIN--   KK-RIDDF-SP  386
Sc  222   IRLDSIQWQTPTIQETKEILKnL[9]PQNDSDGITKK[8]NCELELSQ---TEWYDAKYYLKPSQ-   -VYTYDFF-YP  303
Pp  268   VDLSSIYFKSCKISDIPNVPL--[5]-----------[9]ICEIEELY---TDWTLYEYYHRQE--   KI-KLDHF-QP  332
Sl  509   LDTDTVRWQSLKIESQPDQnDIR[9]PNRDLLAKY--[6]VCEVEDTV---VEWFTEPNIKTKSLK[4]RKLDTEQF-SK  592
Ot  636   IDIDSIEWLDCDTTNPPKE-nVI[9]ASSDKAQKI--[6]ICELEDAV---VEWYDSKYFQRQQ--   EMLNIDFL-NE  712
Sm  867   VDVDKVGWRPLEIFPMQDR----   -----------   ECPIEQEL---FDWYDLDSFQSQSVE   RLQIPDFF-SP  918


Tg  723   CLKHVKTEFSGMDVAVRKSTYRAVRQG-SLGSAACRSWGFPCEI    LPQGVPVVCSLTRWGLQHDR-FLSVQLREGDI  796
Hh  722   CLKHVKTEFSGMDVAVRKSTYRAVRQG-SLGSAACRCWGFPCEI    LPQGVPVVCSLTRWGLQHDR-FLSVQLREGDI  795
Nc  694   HLKHVKTEFSGMDVAVRKSRYRAVRQG-SLGSAACRCWGFPCEI    LPQGLPVVCSLTRRGLQHDR-FLSVQLREGDV  767
Beb 717   HLRHIHTEFFGMDVAVRKSTYRAVRQG-SLGATACRCWGFPCEV    LPQGVPIVCPLTRRGLQHDR-FLSVQLREGDV  790
Cs  719   YLRHFRTEFSGMDVAIRKSTYQAAAPG-SLGVNACRCWGFPCEV    LPAGVPIVCPLTRRGLQHDR-FLSVQLRVGDI  792
Em  1834  ELHHESTEYSGIDLLVRRSLYRAMQEG-TAGAAAQRAWGLQCKV    VSQEAPIVFPLTRKGLLHDR-STCLQLRVGDT 1907
Cc  596   ELHHISTSYSGVDVLVRRSTYNAAREG-SLGAAAKRAWGLQCRV    LPADAPIVFPLTRRGLLHDR-ATSFYLRVGDT  669
To  493   QLVHQVTQVAGIDVITSKSIYVASHTG-MV-VESCNIIGHTVEV    VPKTDSIICMIQRIGLQHDR-ICHVQLRPGDN  565
Bab 459   ELEHQVTQVAGIDVITSRSRFIAAKPG-PV-TEAQNFIGHTMEV    IDARQPVVCMLQRSGVQCDR-LCHVQLRVGDK  531
Pr  525   HLKHINTIYSGIDVTVMKSTYKAIQPG-KLGKKSYNLWGNYFII    EDKNDPVFTFLKREGLQHDYIYHKFYLRVGDY  599
Cm  442   NLIITDTEYAGTDLVTCRVHYKAVKCG--ELSNAEKIFGIDIVV    LPRDYAIICPLKRIGLQHDR-YSPLQIRIDDT  514
Vb  454   HLRHESTVFSGIDIAVSKTRYTAEKPG-RP-EGAERLLGLRLQV    VPRDAAVLVPLTRVGLQHDR-FTNLQVRPGDV  526
Im  276   QLTLKETRCAGVDVITSKLIFQAKNPG--VVSKSYENFGINIKV    LDNKQEAICQARKLGLFQDL-QNEIQLRIGDY  348
Pt  250   HLHLIDCRCAGVDTVAYQFVYEACEIG----SFRNELLGIPIEV    VQQGQEVVTELNKVGLVSDR-ESKLQLRKHDQ  320
Tt  387   YLHLKEVYCAGVDIITSKHIYIAKQPSNGILQKSKEKLGVYIQI    LDDSEDQICEIKRKGLYQDN-ETTIQLRVGDI  461
Sc  304   FLRLMSSEFAGVDVTVSRNTYRAEKIG--IVPDSVARIGILIEI    MPKDEEVTYEVKRMGLVYDR-HRPVELKIGDT  376
Pp  333   YLKLEEVRCSGVDIVTSKHLYKANKVG--QVPKAFENFGVNINI    VPSDKEIIAEVKKIGLFHDL-ENTVELRVGDF  405
Sl  593   QFDISKVEVGGIDFSVWKVQLRAKKNP-LQEYKFYKDLNMGIRI   EKDQSLSIINEIKRSGYLIDK-GQDIELRSGDF  667
Ot  713   LFVVKKYECAGIDGLICKIQLQAVREG----KLSNQYLGISLIV[12]IDQDHGLVNEVKRLGLLIDR-YVDLELRIGDT  795
Sm  919   QLMHMSTEYAGIDVAVSRTRLRAEAQG--VVPQADRILGSRFEV    LPRSSPIILPLKRVGLQHDR-FTKIQLRQGDC  991
```

```
Tg  797   IDYYMSQGGANA*    808
Hh  796   IDYYMSQGGANA*    807
Nc  768   LDYYVSQGGATA*    779
Beb 791   VDYYLSQGGANV*    802
Cs  793   VEFYMSQGGAAV*    804
Em  1908  LESYLTIGGANV*   1919
Cc  670   LDSYLTVGGGTF*    681
To  566   LQFYITKG*        573
Bab 532   VRFYTTKG*        539
Pr  600   VVFYLIKGGNNI*    611
Cm  515   IVLYITKGGAIPE*   527
Vb  527   LYLYISQGGKIIaKR* 541
Im  349   FIFYISKGE*       357
Pt  321   LIFYLTSGD*       329
Tt  462   FVFYISSGD*       470
Sc  377   FVLYISRGG*       385
Pp  406   IIVYISKGD*       414
Sl  668   LLVYVSMGGFEK*    679
Ot  796   LVIYVQRGSS*      805
Sm  992   ILLYISQGGKMT*   1003
```
